# Modeling the MRD state reveals the insomnia of chemotherapy-tolerant persister clones

**DOI:** 10.64898/2026.06.14.732194

**Authors:** Kenta Hata, Taku Sato, Yuki Fukawa, Hiroki Yotsumata, Yumi Mizuguchi, Naozumi Ishimaru, Hiroyuki Harada, Toshiaki Ohteki

**Affiliations:** Department of Biodefense Research, Medical Research Laboratory, Institute of Integrated Research, Institute of Science Tokyo (formerly Medical Research Institute, Tokyo Medical and Dental University (TMDU)), Tokyo 113-8510, Japan; Department of Oral and Maxillofacial Surgical Oncology, Division of Oral Health Sciences, Graduate School of Medical and Dental Sciences, Institute of Science Tokyo, Tokyo 113-8510, Japan; Department of Biochemistry and Molecular Biology, Nippon Medical School Graduate School of Medicine, Tokyo 113-8603, Japan; Department of Oral Pathology, Division of Oral Health Sciences, Graduate School of Medical and Dental Sciences, Institute of Science Tokyo, Tokyo 113-8510, Japan; Division of Surgical Pathology, Institute of Science Tokyo Hospital, Tokyo 113-8510, Japan

## Abstract

Regardless of the success of clinical surgery, disseminated tumor cells (DTCs) can persist in distant organs, with a fraction surviving chemotherapy, which can result in minimal residual disease (MRD), a relevant reservoir for metastatic relapse. Yet, the cellular states that enable the survival and outgrowth of MRD remain poorly defined. Here, using a patient-derived tongue cancer organoid (TCO) model, we recapitulated the key features of chemotherapy-tolerant DTCs by culturing TCOs under growth-factor deprivation and chemotherapeutic stress conditions that mimic the metastatic tissue environment. Clonal-level analyses revealed a distinct subset of cells that retained proliferative capacity without entering a therapy-induced cytostatic state (hereafter referred to as cycling persisters, CPs). CPs exhibited coordinated activation of the IFN signaling pathway, Xenobiotic metabolism, and inflammatory signaling pathways, defining a transcriptional and metabolic program that enables sustained proliferation of CPs under the poor conditions. Consistently, a cell population with similar features was identified in metastatic tissues from patients. Longitudinal clonal tracking demonstrated that early metastatic recurrence is more likely driven by CPs, suggesting a novel mechanism that differs from the prevailing view that relapse arises from reactivation of dormant non-CPs. Our findings highlight a critical therapeutic oversight in relapse prevention.

## Introduction

Despite significant advances in multimodal cancer therapies, recurrence remains a major clinical challenge. Even after seemingly successful treatments, distant relapses can occur in a significant number of patients. This is primarily due to disseminated tumor cells (DTCs), which evade conventional detection methods^1–3^. It is widely considered that a fraction of DTCs survive chemotherapy by entering a dormant, growth-restricted state^4–9^, and metastatic relapse has been attributed to the reawakening of those cells^10–12^. However, direct evidence for such abrupt reactivation in patients remains limited, and it is unclear whether dormant DTCs represent a uniform population with equal relapse-initiating capacities. Rather, DTC populations are likely heterogeneous^9,13,14^, although the specific cellular states that ultimately give rise to clinically manifest metastases remain poorly understood. In this context, DNA damage and mutational processes can occur in non-dividing cells, while the emergence of relapse-initiating clones generally requires subsequent proliferative expansion. How such clones arise from largely growth-restricted populations of DTCs remains an open question, raising fundamental issues about the cellular states and evolutionary dynamics that underlie metastatic recurrence. Thus, defining the cellular state that confers a relapse-initiating capacity on a subset of DTCs is critical for understanding the biological basis of metastatic recurrence and for developing effective prevention strategies.

Using patient-derived tumor organoids (PDTOs), previous studies, including studies from our group, demonstrated that certain PDTO lines exhibit robust drug tolerance not through stable resistant clones but through transient, mutation-independent persister states^15–17^. Here, we established an *in vitro* PDTO platform with AI-based clonal tracking to examine how tumor cells survive and maintain proliferation under minimal-niche conditions after chemotherapy that mimic those in metastatic sites. Our findings show that cell fate is determined early on. Some cells maintain proliferation as cycling persisters (CPs), while others enter growth arrest as non-CPs or die in a stochastic manner. This study proposes an “insomnia” model, distinct from the conventional “long-term dormancy and awakening” model, thereby defining a novel chemotherapy-induced DTC phenotype that contributes to metastatic relapse.

## Results

### A small subset of DTC clones retains proliferative capacity after chemotherapy

To evaluate the proliferative status of DTCs after chemotherapy, we examined cervical lymph nodes, typical organs to which head and neck tumors often metastasize, resected from patients with oral cancer who had received neoadjuvant chemotherapy (NAC) consisting of paclitaxel, carboplatin, and cetuximab (PCE), followed by primary tumor resection and neck dissection (Fig. 1A, B). All identifiable lymph nodes were isolated from the dissected tissues of each of seven patients and the presence of metastatic carcinomas was evaluated by hematoxylin-eosin (HE) staining (Fig. 1A).

**Figure 1.**
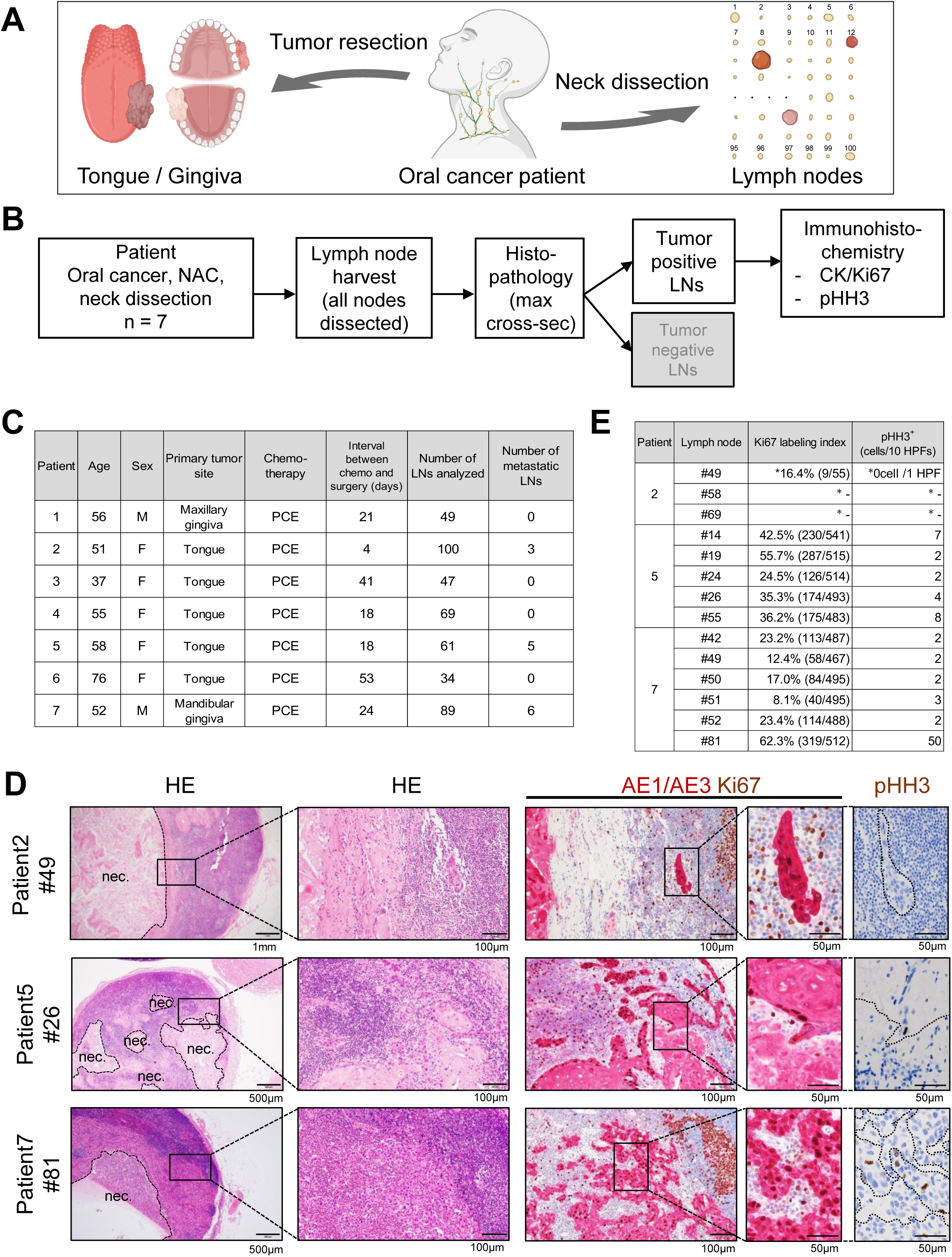
Identification of proliferative tumor clones in metastatic tissues after chemotherapy. **A**, Overview of specimen sampling to assess the status of DTCs. Samples were restricted to cases in which oral cancers were pathologically confirmed as squamous cell carcinomas (SCCs) following neoadjuvant therapy and surgery. The specimens included both primary tumors and lymph nodes. After neck dissection, each lymph node was separated and submitted individually for histopathological examination. Images were created using BioRender.com. **B**, Workflow for the detection and analysis of DTCs. NAC: neoadjuvant chemotherapy, LNs: Lymph nodes. **C**, Summary of clinical samples from patients treated with PCE therapy. PCE: paclitaxel, carboplatin, and cetuximab. **D**, Representative HE staining, double immunostaining for Ki67 and AE1/AE3, and single immunostaining for pHH3 in metastatic lymph nodes after PCE therapy. The dashed lines in pHH3 staining mark the metastatic tumor-stroma boundaries. nec: necrotic area. **E**, Summary of Ki67 labeling index of tumor cells (AE1/AE3 Ki67 double positive cells/total AE1/AE3 positive cells) or the number of pHH3^+^ tumor cells. * Due to extensive necrotic areas, cell counts for these samples were performed only in 1 HPF instead of 10 HPFs. Results of HE staining and immunostaining were reviewed and confirmed by a pathologist. HPF: high power field.

Surgery was performed 4–53 days (median 21 days) after completing NAC (Fig. 1C). Metastatic lesions were detected in three patients, with 34–100 lymph nodes examined per patient (median, 61): Pt. 2 (3/100 nodes), Pt. 5 (5/61 nodes), and Pt. 7 (6/89 nodes) (Fig. 1C).

Most tumor areas in all metastatic lesions exhibited necrosis consistent with chemotherapy-induced cytotoxicity (Fig. 1D, HE). However, Ki67-positive and pan-cytokeratin marker (AE1/AE3)-positive tumor cells were present at the peripheral regions of the tumor nests (Fig. 1D, AE1/AE3 Ki67). Those cells accounted for 8.1%-62.3% (median 23.3%) of viable metastatic carcinoma cells (Fig. 1E). In contrast, only a few mitotically active (pHH3-positive) cells were detected in 10 high-power fields (HPFs) of the tumor tissues in patients 5 and 7, and none were detected in patient 2 (Fig. 1D, pHH3). Notably, one lymph node (#81) from patient 7 contained a higher fraction of pHH3-positive cells (Fig. 1E).

Together, these data suggest that even after systemic chemotherapy, a small subset of metastatic tumor cell clones retains the ability to proliferate in lymph nodes, which could lead to metastatic recurrence.

### *In vitro* modeling of DTCs using patient-derived organoids

Adult epithelial stem cells are maintained within highly specialized, spatially restricted niches that provide growth cues, such as Wnt ligands, epidermal growth factor (EGF) family members, Notch ligands, and bone morphogenetic protein (BMP) modulators, in a concentrated, localized manner^18,19^. Those signals are supplied by defined cellular sources, including specialized epithelial cells and niche-associated stromal components, in a tissue-dependent manner, and are confined to precise microanatomical locations rather than being uniformly distributed throughout the tissue. Consistent with this principle, the normal tongue epithelium proliferated and formed organoids (hereafter referred to as TNOs) in culture medium supplemented with niche factors (NFs), including Noggin, EGF, and R-spondin 1 (hereafter referred to as NF-sufficient medium). Further addition of several factors to the medium, including FGF2, forskolin, Prostaglandin E2, A83-01, nicotinamide, and CHIR99021 (hereafter referred to as TNO-optimal medium), improved the growth of TNOs. In contrast, when NFs were removed (niche-deprived medium), organoids failed to form and did not recover even after exposure to the TNO-optimal medium several days later (Supplementary Fig. 1A–D). These findings were also replicated in normal esophageal squamous epithelial cells (ENOs; Supplementary Fig. 1E, F).

On the other hand, DTCs that extravasate into secondary organs are unlikely to encounter a pre-formed niche. Instead, they lodge in regions with poor niche conditions. Consistent with this, clinical observations and experimental models show that DTCs commonly enter a dormant, non-proliferative, low-metabolic state^6,7,9,20,21^.

Supporting this concept, despite their malignant nature, cancer cells remain dependent on defined niche-derived signals for optimal survival and proliferation. This dependence is exemplified by the requirement for exogenous NFs, such as Wnt ligands, EGF, and other epithelial stem cell-associated signals, for the long-term propagation of PDTOs across multiple epithelial cancer types^15,16,22–26^. Similarly, the withdrawal of niche factors from optimized culture media significantly reduced organoid growth in our TCO models and, in some contexts, induced cell death (Fig. 2A, B). Together, these findings suggest that metastatic tissues do not naturally provide sufficient NFs necessary for robust cancer cell proliferation. Therefore, they are initially unfavorable for early metastatic outgrowth unless supportive components of the niche are subsequently established or remodeled.

**Figure 2.**
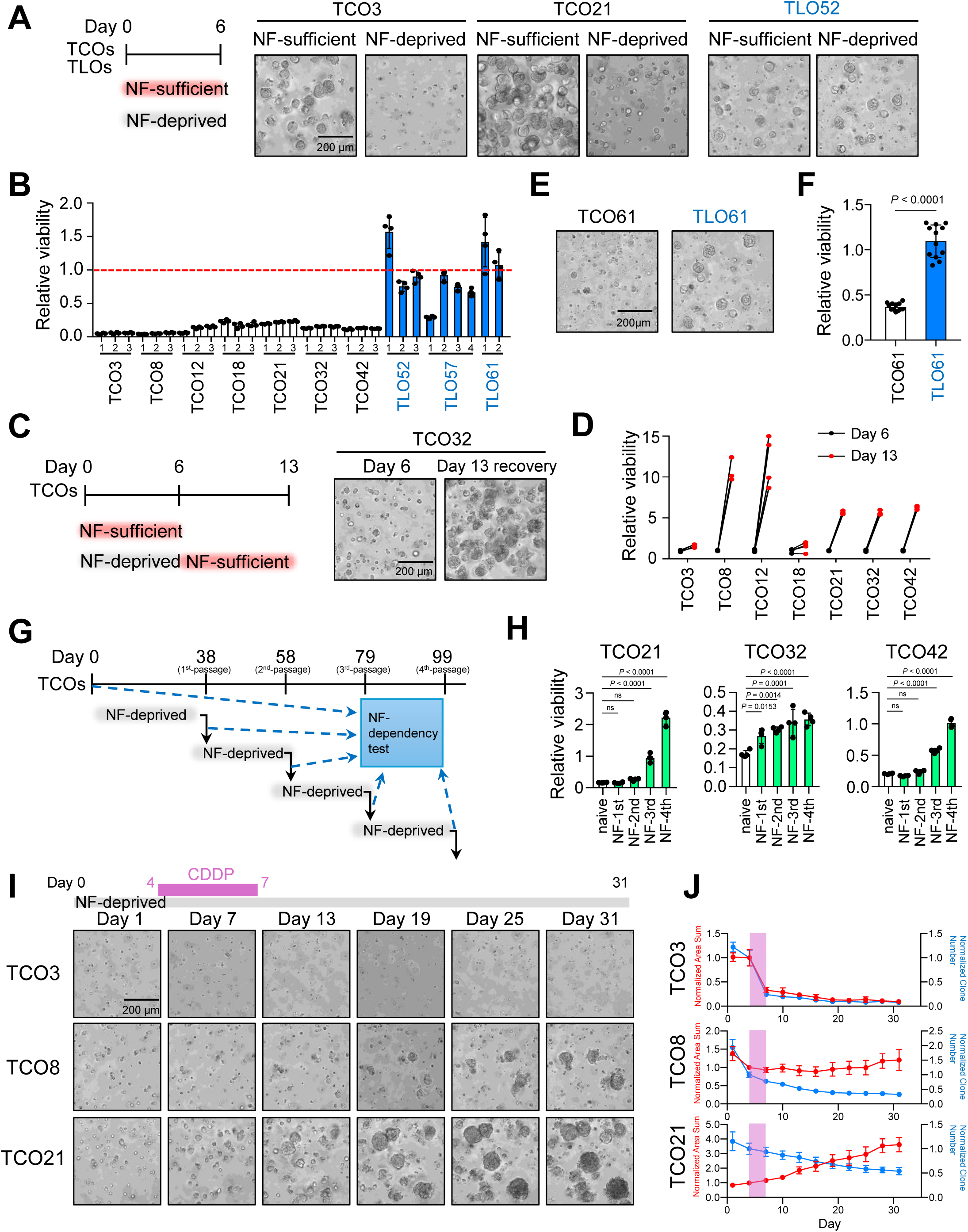
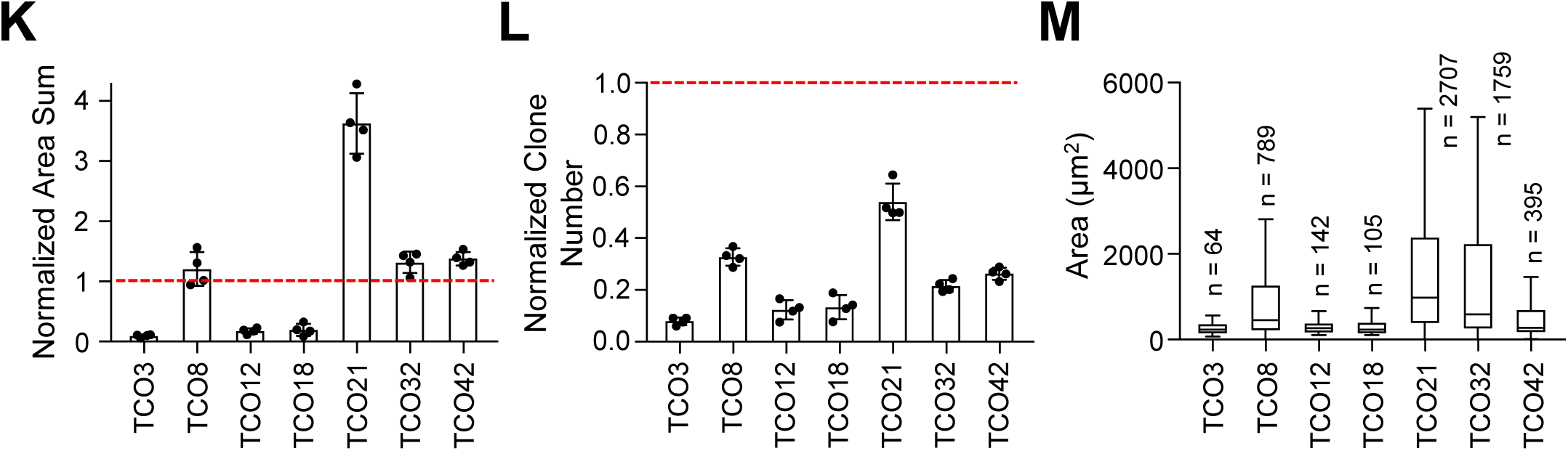
*In vitro* modeling of MRD-like tumor cells using patient-derived organoids. **A**, Schedule for assessing niche factor (NF) dependency in TCOs and TLOs, with representative bright-field images of each culture at day 6. NF: niche factor. Scale bar, 200 μm. **B**, NF dependency in TCOs and TLOs. Relative viability was calculated as the ratio of NF-deprived to NF-sufficient cultures. Data are presented as means ± SD from four independent experiments. **C, D**, Schedule of the transition from NF-deprived to NF-sufficient conditions, with representative bright-field images at day 6 and day 13 (7 days after restoration to NF-sufficient medium) (**C**). Relative viability was calculated for each sample as the ratio of day 13 to day 6 (**D**). Scale bar, 200 μm. **E, F**, Representative bright-field images of paired TCO and TLO derived from the same patient at day 6 (**E**), and NF dependency is shown (**F**). Data are presented as means ± SD of 12 independent experiments. Statistical significance was determined by a two-sided Mann-Whitney U test. Scale bar, 200 μm. **G, H**, Schedule of the long-term culture of primary tumor-derived organoids under NF-deprived conditions. NF dependency was assessed at each passage (**G**). Relative viability was calculated at each time point as the ratio of NF-deprived to NF-sufficient conditions (**H**). Data are presented as means ± SD of four independent experiments. Statistical significance was determined by one-way ANOVA followed by Dunnett’s multiple comparisons test. **I**, Schedule of the culture protocol under NF-deprived conditions with concurrent chemotherapy exposure, with representative bright-field images at days 1, 7, 13, 19, 25, and 31. Scale bar, 200 μm. **J**, Time-course analysis of total clonal area and clone number. Clonal area was quantified from bright-field images and normalized to day 4. The shaded region indicates the period of CDDP treatment. **K, L**, Total clonal area (**K**) and clone number (**L**) at day 31 normalized to day 4. Data are presented as means ± SD from four independent experiments, with individual values shown as symbols. **M**, Box-and-whisker plot showing the distribution of clone areas at day 31. The number of analyzed clones is also indicated.

Notably, while most TCO lines exhibited reduced growth under NF-deprived conditions, some cells appeared to survive. Those cells resumed proliferation when re-exposed to an NF-sufficient medium (Fig. 2C, D). Thus, the capacity of a surviving fraction of cells to withstand NF deprivation without permanently losing their proliferative potential distinguishes tongue cancer cells from their normal epithelial counterparts (Supplementary Fig. 1A–D). Furthermore, TCO organoid lines established from metastatic lymph nodes (hereafter referred to as TLOs) frequently exhibited comparable survival under both NF-deprived and NF-sufficient conditions (Fig. 2B). Similar results were observed when comparing paired TCO and TLO lines derived from the same patient (Fig. 2E, F), demonstrating a substantially reduced dependence of TLOs on exogenous NFs. Consistently, NF-independence increased over the long term in some TCO lines cultured under NF-deprived conditions (Fig. 2G, H). Considering that metastatic lesions are sustained by cancer cell clones derived from primary tumors that have acquired the ability to adapt to microenvironments with poor niches, such as those encountered in lymph nodes, it is plausible that cancer cell clones that survive prolonged culture of primary TCOs under NF-deprived conditions exhibit key functional features of DTCs.

Contrary to the prevailing belief that DTCs remain dormant^4–9^, analysis of post-treatment metastatic specimens revealed that a subset of tumor cells retains proliferative capacity even after chemotherapy (Fig. 1D, E). We therefore modeled this clinically relevant state *in vitro* by culturing tumor cells under NF–deprived and chemotherapeutic conditions that mimic at least in part the niche-poor, treatment-exposed metastatic microenvironment. First, we examined the impact of NF-deprived conditions on the response to cisplatin (CDDP), a standard chemotherapeutic agent for oral cancer. IC50 values were calculated from the dose-response curves for each TCO after 10 days of culture in NF-sufficient medium or in NF-deprived medium, and CDDP was administered from day 4 to day 7. Notably, all TCOs showed a marked increase in IC50 when cultured in NF-deprived medium compared to NF-sufficient medium (Supplementary Fig. 2A), which suggests that DTCs residing in an NF-deprived environment display enhanced resistance to chemotherapy. A similar increase in chemotherapeutic resistance under these culture conditions was also observed in an esophageal squamous cell carcinoma (ESCC) model (Supplementary Fig. 2B).

To monitor dynamic changes in individual surviving cancer cell clones, conventional viability assays based on bulk ATP measurements, such as CellTiter-Glo, have a couple of limitations. Since the assay requires reagent-induced cell lysis, continuous long-term tracking of the same well is not feasible. In addition, these assays provide only bulk measurements of cell viability and therefore do not permit assessment at the single-clone level. To address those issues, we developed an image-based analytical framework employing time-resolved, high-resolution imaging of 3D organoid cultures. Within that framework, we identified and quantitatively assessed viable cell clusters based on their projected areas. We employed a deep learning–based artificial intelligence (AI) approach for cell viability classification. Specifically, we used paired datasets consisting of brightfield images and corresponding propidium iodide (PI) stained images from the same fields for iterative model training. Through this process, we developed a model that can accurately identify dead cells using brightfield images alone (Supplementary Fig. 3). Dead cells identified by PI staining were used as ground truth (Supplementary Fig. 3A, B), and deep learning models were iteratively trained for the precise detection of dead cells (Supplementary Fig. 3C, D). Consequently, the identification of dead clones achieved more than 95% accuracy, with the area sum and clone number as key metrics (Supplementary Fig. 3E, F). In this method, excluding dead-cell regions enabled us to define viable cells. The number of viable cells estimated by this method was highly correlated with measurements obtained by CellTiter-Glo (Supplementary Fig. 3G). Taken together, these observations indicate that our image-based analytical framework is well-suited for longitudinal tracking of individual clones within the same well over extended culture periods.

Using this method, organoids were monitored for up to 1 month in NF-deprived medium. CDDP was administered from day 4 to day 7 of the culture at a concentration corresponding to the maximum plasma concentration (C_max_) observed in patients. Specific organoid lines, such as TCO8, TCO21, TCO32, and TCO42, were found to contain a relatively higher proportion of cancer cell clones that continued to proliferate without entering a cytostatic state (Fig. 2I-M, Supplementary Fig. 4A, B), implying the presence of proliferative cancer cell clones under poor-niche and chemotherapeutic conditions in patients as a potential source of tumor recurrence. These clones are referred to as CPs. In contrast, proliferating cancer cell clones were only rarely observed in organoid lines such as TCO3, TCO12, and TCO18 under the same culture conditions (Fig. 2I–M, Supplementary Fig. 4A, B). Furthermore, following the extended culture under NF-deprived conditions combined with chemotherapy exposure, subsequent restoration of NF-sufficient conditions failed to induce re-expansion of previously non-proliferative clones (Supplementary Fig. 4C, D). A similar inter-organoid line heterogeneity was also observed in ESCCOs (Supplementary Fig. 4E, F). The frequency of CPs was highly reproducible across independent experiments, indicating that the underlying determinants of this phenotype are likely driven by stable genomic and/or epigenomic alterations maintained during serial passaging.

### Clonal-level analysis of 3D cultures defines the heterogeneity of chemotherapy responses

Although a fraction of cells at metastatic sites retain the ability to proliferate even after systemic chemotherapy (Fig. 1D, E), snapshot analyses of pathological specimens cannot determine the subsequent fate of these cells. To directly interrogate the long-term fate of CPs versus non-CPs following exposure to chemotherapeutic drugs, we developed a novel *in vitro* clonal-tracking system that enables the precise monitoring of individual tumor clones. This system allows us to study their distinct proliferative behaviors and contributions to relapse. In this system, after AI-assisted segmentation (Supplementary Fig. 3A-D), individual clonal aggregates were labeled, and their centroids were computationally registered across serial images. This alignment procedure reliably tracked the same clones throughout the entire observation period (see Methods). Representative outputs are shown in Fig. 3A, demonstrating accurate and stable clonal identification over time.

**Figure 3.**
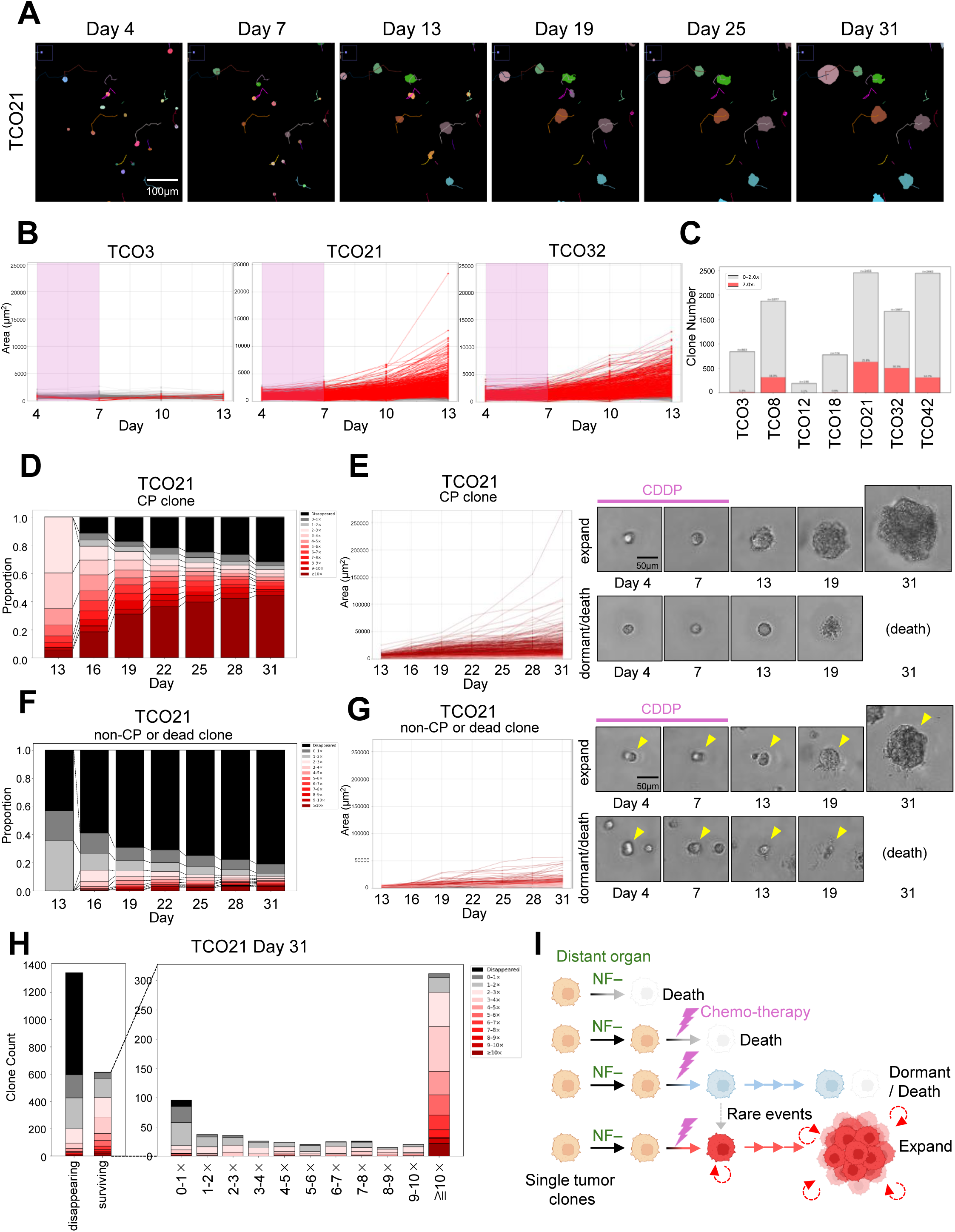
Clonal behavior analysis of tumor cells reveals heterogeneity in chemotherapy response in 3D *in vitro* cultures. **A**, Representative AI-annotated images of clone tracking in TCO21. Clonal areas from day 4 (start point) to day 31 (end point) are shown with their trajectories. Scale bar, 100 μm. **B**, Changes in clonal areas of individual clones from day 4 to day 13, quantified from bright-field images. Red lines indicate CP clones (≥ 2-fold expansion in area between day 7 to day 13), whereas gray colored lines indicate non-CP or dead clones (< 2-fold change). The shaded regions indicate the period of CDDP treatment. **C**, Stacked bar graph showing the proportions of CP and non-CP/dead clones. Percentages were calculated relative to the total number of clones at day 4. **D**, 100% stacked bar graph showing the distribution of fold changes in clonal area for CP clones of TCO21 at each time point (days 13-31). Fold changes were calculated for each clone at each time point relative to day 7 and were categorized into discrete ranges, with increasing values shown as progressively darker shades of red (≥ 10-fold as the darkest), 1-2-fold in light gray, and 0-1-fold in dark gray, and disappeared clones in black. Notably, only clones exhibiting ≥ 10-fold expansion relative to day 7 progressively become dominant over time, suggesting that a limited subset of clones possesses sustained proliferative capacity and may ultimately contribute to tumor outgrowth. **E**, Left, changes in clonal area of individual CP clones from day 13 to day 31 (left). The color of each line corresponds to the categories shown in Fig. 3D. Right, representative bright-field images of an expanding CP clone (upper right) and a dormant or dying CP clone (lower right) are shown. Scale bar, 50 μm. **F**, 100% stacked bar graph showing the distribution of fold changes in clonal areas for non-CP or dead clones of TCO21 at each time point (days 13-31). Fold changes were calculated for each clone at each time point relative to day 7 and were categorized into discrete ranges, with increasing values shown as progressively darker shades of red (≥ 10-fold as the darkest), 1-2-fold in light gray, and 0-1-fold in dark gray, and disappeared clones in black. Notably, few non-CP clones at day 13 become expanding clones over time. **G**, Left, changes in clonal areas of individual non-CP or dead clones from day 13 to day 31 (left). The color of each line corresponds to the categories shown in Fig. 3F. Representative bright-field images of an expanding non-CP clone (upper right) and a dormant or dying non-CP clone (lower right) are shown. The arrowheads indicate the tracked clones. Scale bar, 50 μm. **H**, Tracked clones were first categorized as disappearing or surviving by day 31 (left). From another perspective, surviving clones were further stratified based on changes in clonal area at day 31 relative to day 7 (right). Stacked bar graphs show changes in clonal areas of individual clones at day 13 relative to day 7. The color of each segment corresponds to the categories shown in Fig. 3D, F. Notably, the most prominently expanding clones also exhibited rapid expansion between day 7 and day 13, indicating that clonal fate is largely determined during chemotherapy exposure rather than during the subsequent dormant survival phase. **I**, Based on clonal tracking, chemotherapy-exposed disseminated tumor clones in distant organs can be classified into four fate types. In NF-deprived metastatic niches, clones may die, may survive but remain sensitive to additional chemotherapy-induced death, may survive chemotherapy but subsequently undergo long-term dormancy or death under prolonged NF deprivation, or may survive chemotherapy and maintain sustained proliferative capacity. In contrast, the reawakening of deeply dormant clones into expanding clones was rarely observed. Images were created using BioRender.com.

Using this system, we monitored thousands of individual clones immediately after exposure to chemotherapeutic agents across multiple TCO lines, including TCO3, TCO8, TCO12, TCO18, TCO21, TCO32, and TCO42. Changes in two-dimensional clonal areas were quantified based on microscopic images from day 4 (treatment initiation) to day 13 (Fig. 3B, Supplementary Fig. 5A,B). Clones whose area increased by more than twofold from day 7 to day 13 were defined as CPs and are highlighted in red. In contrast, clones exhibiting less than a twofold increase were classified as non-CPs or dead (gray). The frequency of CP formation varied markedly between TCO lines: TCO8, TCO21, TCO32, and TCO42 generated relatively abundant CPs (hereafter referred to as CP-rich TCOs), whereas TCO lines TCO3, TCO12, and TCO18 hardly produced CPs (hereafter referred to as CP-poor TCOs) (Fig. 3C).

Next, we tracked the long-term fate of CPs in CP-rich TCO lines from day 13 to day 31. Notably, clonal behaviors became increasingly polarized over time. By day 31, the surviving clones were primarily composed of cells that had either expanded minimally or grown more than tenfold compared to their size on day 7. These results imply that the proliferative capacity of CPs that emerge after chemotherapy exposure is heterogeneous. Not all CP clones maintain continuous growth, and only a subset of them can evade growth arrest, retaining the ability to proliferate and form large aggregates over time (Fig. 3D, E, Supplementary Fig. 6, 8A, B). In addition, we observed that non-CP clones in CP-rich TCOs began to proliferate slowly from day 13 onward under NF-deprived conditions, a phenomenon we defined as “slow-awakening”, characterized by a greater than twofold increase in area relative to day 7. “Slow-awakening” clones were only rarely observed in CP-rich TCOs (Fig. 3F, G, Supplementary Fig. 8C, D) and were negligible in CP-poor TCOs (Supplementary Fig. 7A-C). Finally, we classified clones as “disappearing” or “surviving” at day 31 and quantified the extent of expansion among surviving clones between day 13 and day 31. Notably, among clones that expanded by more than tenfold, the vast majority had already increased in size by at least twofold between day 7 and day 13 (Fig. 3H). This trend was consistently observed across other CP-rich TCOs (Supplementary Fig. 6).

Together, our data suggest that CPs arise from pre-existing clones whose molecular state enables their survival without entering a cytostatic, dormant program. Those clones maintain the capacity to proliferate under cytotoxic stress, challenging the conventional “long-term dormancy and awakening” model by demonstrating that persistent proliferation, rather than dormancy, underlies long-term survival (Fig. 3I).

### CPs exhibit MRD features of chemotherapy-treated patients

To elucidate the molecular basis of the emergence of chemo-resistant CPs under NF-deprived conditions, cancer cells derived from CP-rich or CP-poor TCO lines were either exposed to chemotherapeutic agents under NF-deprived conditions or were maintained under drug-free, NF-sufficient conditions, and interline differences in gene expression profiles were analyzed. Unsupervised principal component analysis (PCA) revealed that samples clustered primarily by organoid line rather than separating into distinct groups based on CP-rich versus CP-poor status and culture conditions, including NF-sufficient versus NF-deprived with chemotherapeutic treatment (Fig. 4A).

**Figure 4.**
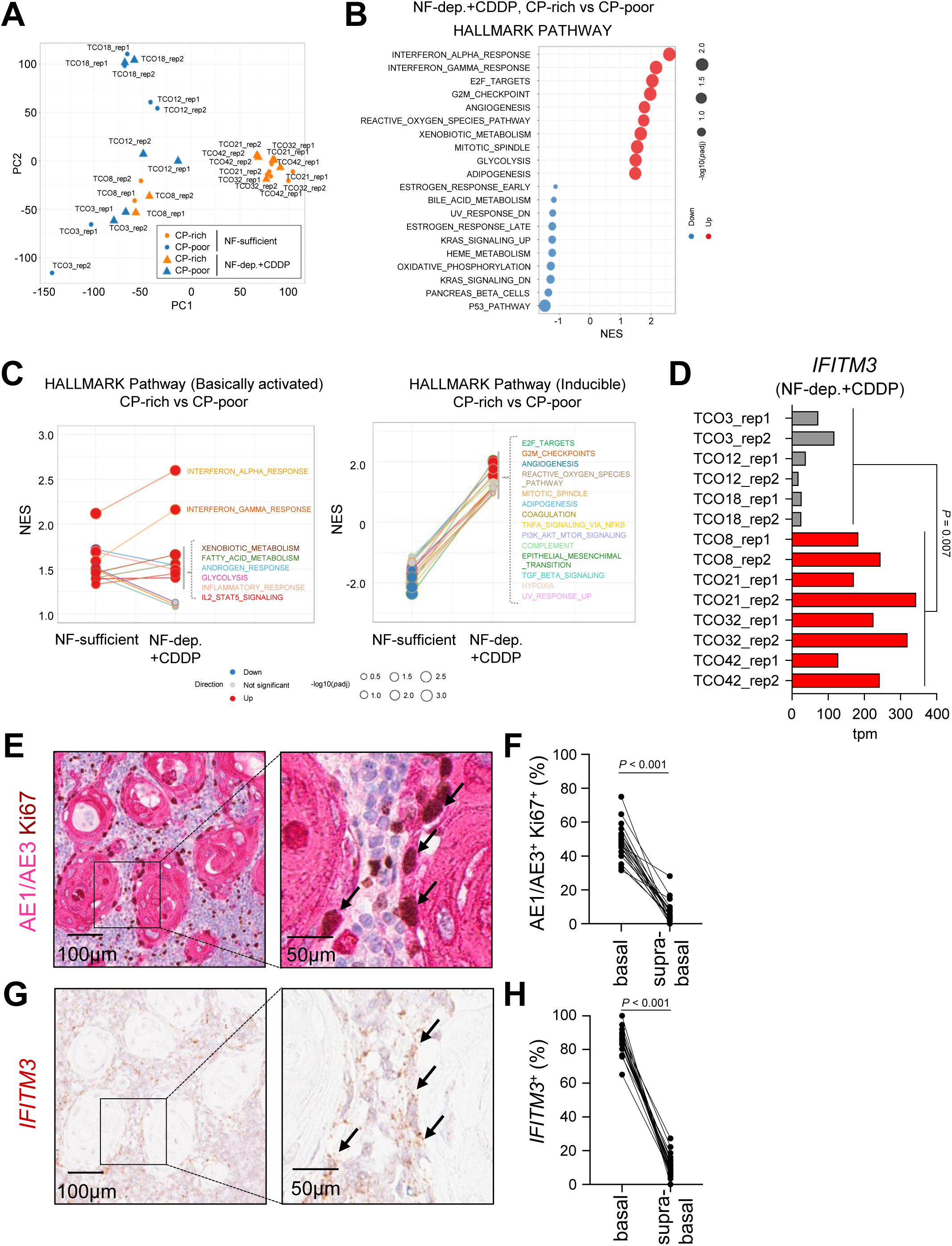
CP phenotype recapitulates MRD-like features in chemotherapy-treated metastatic lesions. **A**, PCA plots showing gene expression variations among CP-rich (TCO8, TCO21, TCO32, TCO42) and CP-poor (TCO3, TCO12, TCO18) lines. Cultures under NF-sufficient conditions are represented by circles, whereas those under NF-deprived conditions with CDDP exposure are represented by triangles. Cells under NF-sufficient conditions were harvested on day 7. Cells under NF-deprived conditions were treated with CDDP from day 4 to day 7 and were harvested on day 7. **B**, GSEA comparing transcriptional profiles between CP-rich and CP-poor lines under NF-deprived conditions with CDDP treatment, using the same culture conditions as described in **A**. The top 10 HALLMARK gene sets (MSigDB) significantly enriched in either CP-rich (red) or CP-poor (blue) lines (NOM p-value < 0.05) are shown. **C**, GSEA was performed for CP-rich and CP-poor lines under NF-sufficient or NF-deprived conditions with concurrent CDDP exposure. Differentially enriched pathways between the two groups were identified for each condition and plotted, and their changes across conditions were assessed. Pathways consistently enriched in CP-rich lines (left) and those specifically enriched under NF-deprived conditions with concurrent CDDP exposure (right) are shown. **D**, Comparison of *IFITM3* mRNA expression between CP-rich (red bars) and CP-poor TCO lines (grey bars). TPM values are shown. Statistical significance was determined using a two-sided Mann–Whitney test. **E-H**, Representative double immunostaining for Ki67 and AE1/AE3 (**E**), and RNAscope-based detection of *IFITM3* mRNA (**G**) in serial pathological sections from Patient 5 shown in Fig. 1C. Corresponding cells positive for both markers in serial sections are indicated by arrows. The frequencies of Ki67- and AE1/AE3-double positive proliferating tumor cells (**F**), as well as *IFITM3*-expressing cells (**H**), were evaluated in the basal and supra-basal layers of the lymph node tumor nests. Results of immunostaining and RNAscope were reviewed and confirmed by a pathologist. Statistical significance was determined by two-sided Wilcoxon test. Scale bars, 100 μm (left panels of **E** and **G**), 50 μm (right panels of **E** and **G**).

Next, we compared the gene expression profiles of cancer cells derived from CP-rich or from CP-poor lines under NF-deprived conditions in the presence of chemotherapeutic agents. Gene set enrichment analysis (GSEA) revealed that the most significantly upregulated pathways in CP-rich lines, compared to CP-poor lines, included the INTERFERON_ALPHA_RESPONSE and INTERFERON_GAMMA_RESPONSE pathways. In addition, the E2F_TARGETS pathway, which reflects proliferative activity, was activated, whereas genes associated with the P53_PATHWAY, which is implicated in growth arrest and cell death following chemotherapy, were suppressed. These findings indicate that CPs retain their proliferative capacity even under highly stringent culture conditions. Consistently, the activation of genes related to cell cycle checkpoints (G2M_CHECKPOINT) was also observed. Moreover, multiple metabolic pathways, including the REACTIVE_OXYGEN_SPECIES_PATHWAY, XENOBIOTIC_METABOLISM, GLYCOLYSIS, and ADIPOGENESIS, were upregulated, which suggests that transcriptional metabolic adaptation is a key feature underlying CP emergence across different lines (Fig. 4B). We also compared the gene expression profiles of CP-rich and CP-poor lines before and after CDDP exposure. Several pathways, including the IFN SIGNAL PATHWAYS, XENOBIOTIC_METABOLISM, GLYCOLYSIS, FATTY_ACID_METABOLISM, and INFLAMMATORY_RESPONSE, remained consistently active in CP-rich lines regardless of CDDP treatment. In addition, compared to CP-poor lines, the following pathways were relatively weak before treatment but were more strongly induced after drug exposure in CP-rich lines: cell proliferation (E2F_TARGET, G2M_CHECKPOINTS, MITOTIC_SPINDLE), inflammatory signaling (COAGULATION, TNFA_SIGNALING_VIA_NFKB, COMPLEMENT), and epithelial–mesenchymal transition (EPITHELIAL_MESENCHYMAL_TRANSITION, TGFB_SIGNALING). The activation of these pathways suggests their involvement in establishing a cellular state permissive for CP emergence (Fig. 4C). Previous studies have reported that cancer cell clones surviving chemotherapy adopt a diapause-like metabolically dormant state^27–30^. In contrast, CPs observed in our model exhibited active transcriptional metabolic adaptation, indicating that they do not conform to the previously proposed model in which low metabolic activity enables evasion of chemotherapeutic effects. Furthermore, our findings suggest that pre-existing transcriptional states facilitate the rapid induction of adaptive responses required for CP emergence.

Minimal residual disease (MRD) is inherently difficult to detect in patients by conventional clinical evaluation^1–3^. However, detailed analysis of metastatic lymph nodes resected after NAC revealed viable cancer cells, a subset of which clearly retained proliferative capacity (Fig. 1D, E). Those cells correspond to CPs and suggest that similar populations may persist across multiple organs in patients. Within the heterogeneous MRD compartment, such cells could represent a subpopulation with the potential to drive disease recurrence.

To determine whether *in vitro*-induced CPs recapitulate the MRD state *in vivo*, we next performed marker-based validation. Among the IFN signaling–related genes markedly upregulated in CP-rich lines (Fig. 4B, C), we selected *IFITM3* as a representative marker (Fig. 4D, Supplementary Fig. 9, red arrowheads) and examined its expression in cancer cells that had metastasized to lymph nodes in Pt. 5. Ki-67^+^ proliferative tumor cells were predominantly localized to the basal compartment of the tumor nests, whereas differentiated tumor cells in the supra-basal keratinizing layers exhibited little to no proliferative activity (Fig. 4E, F, Supplementary Fig. 10A, B). Of note, RNA-scope analysis demonstrated that *IFITM3* mRNA was strongly expressed in Ki-67^+^ proliferative, but not in Ki-67^-^non-proliferative, tumor cells within basal regions (Fig. 4G, H, Supplementary Fig. 10C, D), suggesting that *IFITM3* is a novel marker for detecting CPs in metastatic lymph nodes. Collectively, our *in vitro* MRD model recapitulated MRD features observed in patient metastatic tissues, which were concentrated in CPs, a relevant source of recurrence.

We further evaluated the validity of our *in vitro* MRD model using previously reported mouse metastasis models. Aouad et al. characterized the heterogeneity of DTCs in a mouse breast cancer metastasis model and identified two distinct populations based on proliferative status: active and inactive DTCs^21^. Active DTCs were characterized by activation of cell cycle–related pathways, including E2F_TARGET and G2M_CHECKPOINT, whereas inactive DTCs exhibited reduced activity of those pathways and a low-proliferative state. Just as with the characteristics of CPs in our MRD model, active DTCs were enriched for multiple biological pathways, including GLYCOLYSIS, REACTIVE_OXYGEN_SPECIES_PATHWAY, XENOBIOTIC_METABOLISM, INTERFERON_GAMMA_RESPONSE, ADIPOGENESIS, and COAGULATION (Supplementary Fig. 11A). Zhang et al. demonstrated that DTCs residing in the lungs exist in a dormant state, whereas bleomycin treatment triggers their awakening and acquisition of proliferative capacity^31^. We re-analyzed the published dataset, which revealed significant enrichment of multiple pathways, including INTERFERON_GAMMA_RESPONSE, INTERFERON_ALPHA_RESPONSE, INFLAMMATORY_RESPONSE, TNFA_SIGNALING, IL2–STAT5_SIGNALING, XENOBIOTIC_METABOLISM, and ADIPOGENESIS. Again, these transcriptional features closely mirror those observed in our CP population (Supplementary Fig. 11B).

In summary, our “insomnia” model demonstrates that CPs that emerge from the beginning of culture conditions mimicking the harsh microenvironment of metastatic tissues reproduce the key characteristics of metastatic CPs found in the MRD of the patients. In addition, the “insomniac” DTCs (CPs) share a transcriptional program with late-awakening DTCs in the conventional “long-term dormancy and awakening” model, highlighting the validity and utility of our model for identifying therapeutic strategies targeting early versus late relapse-initiating cell populations.

## Discussion

In this study, we established culture conditions that recapitulate the metastatic tumor microenvironment and applied them to a diverse panel of TCO lines. Under these conditions, we identified a population of CPs that survive chemotherapy while retaining their proliferative capacity. We further characterized these CPs based on their gene expression profiles. In addition, we identified CP-associated gene markers and examined their spatial expression patterns in patient-derived metastatic tissues, demonstrating the presence of a similar cell population in metastases from chemotherapy-treated patients. These cells are likely to represent a population capable of contributing to early disease recurrence. Collectively, our model provides a valuable platform for investigating the molecular basis of cancer recurrence.

Our findings are not readily reconciled with a simple long-term dormancy-and-awakening model, in which DTCs persist in a deeply quiescent state and subsequently re-enter the cell cycle to drive metastatic outgrowth^10–12,32^. Instead, our results suggest that the proliferative potential of individual clones is largely determined prior to chemotherapy exposure, and that treatment-induced damage dictates whether clones with proliferative potential accelerate their expansion (insomnia) or clones lacking the potential enter a durable, non-expanding state. Following extended culture (1 month) under NF-deprived conditions combined with chemotherapy exposure, subsequent restoration of NF-sufficient conditions failed to induce robust re-expansion of previously non-proliferative clones (Supplementary Fig. 4C, D). These observations underscore the importance of the “insomnia” model in the post-treatment dynamics of tumor cells and suggest that, at least under these conditions, the conventional “long-term dormancy and awakening” model is rare.

At the same time, several limitations of our study should be considered. Organoid culture systems are inherently constrained in duration and do not fully recapitulate the long-term persistence of DTCs in patients. It therefore remains possible that prolonged survival within native microenvironments could induce additional cellular adaptations that enable delayed reawakening, as proposed in the long-term dormancy model.

Despite the success of clinical surgery and chemoradiotherapy, DTCs persist as MRD in tissues distant from the primary tumor site. The suppression of recurrence could be achieved by keeping MRD in a sleeping state. Given that this represents a key concept for future cancer control strategies, our model will serve as a vital tool for elucidating the mechanisms underlying CPs and for developing control methods based on those mechanisms, thereby helping prevent recurrence.

## Methods

### Human tissue samples

The pathological specimens used to detect DTCs were obtained from samples collected at the Institute of Science Tokyo Hospital. Samples were restricted to cases in which oral cancer patients underwent surgery after receiving PCE neoadjuvant therapy and had a pathologically confirmed diagnosis of squamous cell carcinoma (SCC).

Patient samples used to establish organoids were obtained from the Institute of Science Tokyo Hospital, and the Jichi Medical University Hospital. None of the patients had received NAC. Normal, lymph node, and tumor tissues were obtained from surgically resected specimens. Tumor tissues were collected from non-necrotic regions. Lymph node samples were obtained from clinically apparent metastatic lymph nodes, including submandibular, submental, and deep cervical (superior and middle) nodes. Subsequent histopathological examination confirmed SCC in the primary tumor and metastasis to the lymph nodes. The study was approved by the institutional ethical committees of both institutes. Written informed consent was obtained from all patients, and tissues were used solely for research purposes. A previously established library of normal epithelium and cancer organoids generated in our laboratory was also used.

### Organoid establishment

Tissues were cut into small pieces and digested with 0.125% or 0.025% trypsin for 30 min at 37 ℃ with vigorous pipetting every 10 min. The reaction was quenched by adding PBS lacking Ca^2+^ and Mg^2+^ supplemented with 10% FCS (hereafter, 10% FCS/PBS), and the suspension was passed through a 100 µm cell strainer (Corning). Cells were then centrifuged at 400 × *g* for 5 min, and the pellets were resuspended in 1 mL Advanced DMEM/F-12 medium (Thermo Fisher Scientific) containing Primocin (Invivogen), 1 mM HEPES (Nacalai), and L-alanyl-L-glutamine (Nacalai) (hereafter, AdDF+++). After centrifugation, the pellets were resuspended in Matrigel (Corning) and dispensed into culture plates. The plates were incubated at 37 °C for 20 min to allow the Matrigel to solidify, after which culture medium was added to each well. For culturing cancer organoids, AdDF+++ medium supplemented with 1×B27 (Invitrogen), 1.25 mM N-acetylcysteine (Sigma-Aldrich), 10% RspoI-culture supernatant of the 293T-HA-RspoI-Fc cell line (provided by Calvin Kuo of Stanford University), 100 ng/mL Noggin (Miltenyi), 50 ng/mL EGF (Peprotech), and 10 µM Y-27632 (Nacalai), which is collectively referred to as NF-sufficient medium, was added to each well. In addition to these components, the medium for normal organoids was supplemented with 5 ng/mL FGF2 (PeproTech), 1 µM forskolin (Tocris), 1 µM Prostaglandin E2 (Tocris), 5 nM A83-01 (Nacalai), 10 mM nicotinamide (Sigma), and 0.3 µM CHIR99021 (Chemscene), which is collectively referred to as TNO-optimal medium. Cells were cultured at 37 ℃ with 5% CO_2_, and the medium was changed every 3-4 days. For passaging, organoids were collected, treated with TrypLE Express (Thermo Fisher Scientific) at 37 ℃ for 15 min, and mechanically dissociated by pipetting. The cells were then washed with 10% FCS/PBS and centrifuged at 440 × *g* for 5 min. The cell pellets were resuspended in AdDF+++ and centrifuged again at 440 × *g* for 5 min, then subsequently resuspended in Matrigel. Organoids were passaged every 7-14 days.

### Histology and immunostaining

Human tissues were fixed with 10% phosphate-buffered formalin (pH 7.4, Fujifilm Wako Chemicals) and were prepared for histological examination. Formalin-fixed, paraffin-embedded (FFPE) tissue blocks were cut into 3 μm-thick sections for HE staining. Only lymph nodes with histologically confirmed SCC metastases following PCE neoadjuvant therapy were selected for the immunohistochemical analysis. FFPE tissue blocks were cut into 3 μm-thick sections for immunohistochemical analysis. Both double staining for Ki-67 and AE1/AE3, and single staining for phosphor-histone H3 (pHH3) were performed.

Double immunohistochemical staining for Ki-67 and AE1/AE3 was performed using the BOND-III system (Leica Biosystems). Antigen retrieval for both markers utilized BOND Epitope Retrieval Solution 2 at 100°C for 20 min. Sequential staining was performed starting with anti-Ki-67 (clone MIB-1, 1:200; Agilent Technologies) detected via the BOND Polymer Refine Detection (DAB) system, followed by anti-AE1/AE3 (1:4; Agilent Technologies) detected utilizing the BOND Polymer Refine Red Detection (Alkaline Phosphatase) system. For pHH3 single staining, a manual staining protocol was performed. Antigen retrieval was conducted using a microwave at 97°C for 40 min in Histofine pH 9 buffer (Nichirei Biosciences). Sections were incubated with an anti-pHH3 primary antibody (1:500 dilution; Cell Marque), followed by detection using the Dako EnVision+ System-HRP Labeled Polymer Anti-Rabbit (Agilent Technologies). pHH3 positivity was evaluated by a pathologist based on previously published criteria^33^.

### Cell viability assay

For testing NF dependency, organoids derived from primary tumors, metastatic lymph nodes, or normal epithelium were dissociated into single cells, embedded in Matrigel at 2000 cells/10 µL, and seeded into 96-well plates. After Matrigel solidification, NF-sufficient medium (AdDF+++ supplemented with B27, N-acetylcysteine, 10% RspoI-culture supernatant of the 293T-HA-RspoI-Fc cell line, Noggin, and EGF) or NF-deprived medium (AdDF+++ supplemented only with B27 and N-acetylcysteine) was added. Y-27632 was included immediately after passaging and was removed at the next medium change. Cell viability was quantified on day 6 using CellTiter-Glo (Promega). In some experiments, NF-deprived medium was replaced with NF-sufficient medium on day 6, and cells were cultured for an additional 7 days to assess survival after NF-deprivation. CellTiter-Glo was performed on day 13.

For CDDP dose–response curve analysis, dissociated tumor cells were embedded in Matrigel at 2000 cells/10 µL and seeded into 96-well plates (10 µL per well). Cells were treated with various concentrations of CDDP (Fujifilm Wako Chemicals) from day 4 to day 7 after seeding. Cell viability was measured on day 10 using CellTiter-Glo (Promega).

For long-term NF-deprived cultures, dissociated tumor cells were embedded in Matrigel and then cultured in NF-deprived medium as described above. CDDP was added from day 4 to day 7, and Y-27632 was included immediately after passaging. Cultures were maintained until day 31 with medium changes every 3 days.

### Microscopic imaging and deep-learning

Microscopic images were acquired using Cell3iMager NX (SCREEN). To enable longitudinal quantification of viable organoid clone areas (defined as areas of cell clusters detected in microscopic images) without sacrificing the wells, a deep-learning-based image analysis approach was employed. Organoid clones were first annotated in bright-field images by manual labeling. These annotations were then used to train a deep learning model to extract organoid clones irrespective of their viability status, and a trained model was generated for subsequent analysis. Next, based on the model file, PI staining was used to annotate dead (PI-positive) and viable (PI-negative) cells. The annotated dataset was then used to train a deep learning model to discriminate between viable and dead cells solely from bright-field images. In the initial output, the AI classified the clones into four groups based on their positive/negative status for PI staining and the results of AI initial predictions. Type I: clones labeled as “alive” with PI fluorescence intensity values < 20, Type II: clones labeled as “dead” with PI fluorescence intensity values ≥ 10, Type III: clones labeled as “dead” despite PI fluorescence intensity values < 10, Type IV: clones labeled as “alive” despite PI fluorescence intensity values ≥ 20. Types I and II were excluded from additional training. In contrast, Type III clones were retrained as “alive” for additional training. In particular, considering that expanding organoids naturally incorporate internal dead cells even within viable clusters^34,35^, clones labeled as Type IV were generally categorized as “alive”, while morphologically obvious dead cells and debris were manually re-annotated. These deep-learning models were run repeatedly until significant correlations were observed between PI-positive clones predicted as “Dead” (Type II) and all clones predicted as “Dead” (Types II and III) both for area and for clone count. In addition, we verified that the average accuracy for area and clone count reached to over 95%. To minimize noise, debris and objects smaller than 100 µm^2^ were excluded from the clone recognition. To detect MRD clones, sequential clone imaging was performed every 3 days from day 1 to day 31 after seeding.

### Clone tracking

Image stacks (serial 2D fields or time series) were aligned using the manual landmark-based alignment function of the TrakEM2^36^ plugin in Fiji^37,38^ with a rigid transformation model. For each pair of consecutive frames, three stationary debris particles within the field of view were manually annotated as fiducial landmarks to guide the alignment. Organoid foreground was segmented using the Color Threshold tool in HSB space, with empirically tuned thresholds (Red channel). Resulting masks were converted to 8-bit binary images. Spurious foreground was removed in Fiji, and organoid-sized objects were extracted using size- and shape-based filtering (Analyze Particles), and manual correlations were applied when needed. Binary mask images were converted into labeled images using Connected Components Labeling with MorphoLibJ^39^. These labeled objects were then tracked with

TrackMate^40,41^ using the Label Image Detector and the LAP tracker (frame-to-frame linking max distance: 30 px; track segment gap-closing max distance: 50 px; max frame gap: 2; track segment splitting: 30 px; track segment merging: 20 px). Spot data for each cell cluster were curated, and the resulting table included TrackID (which is a unique ID associated to each individual clone for tracking purposes), Time, and Area (px^2^). Rare fusion (≤ 1.6%) or splitting (≤ 2.0%) events (Supplementary Fig. 5B) were corrected by aggregating the areas of all TrackIDs involved. This aggregated area was treated as a single clone and used consistently for area quantification at the time points immediately preceding and following the event.

All data processing and statistical analyses were performed in Python (v3.13). Graphical outputs were generated using matplotlib.

### Bulk RNA sequencing and data analysis

RNA-seq library preparation, sequencing, mapping, and gene expression analyses were performed by CyberomiX Inc.. Double-stranded cDNA was synthesized directly from 1000 cells according to the manufacturer’s instructions using a SMART-Seq v4 Ultra Low Input RNA Kit (Takara Bio). Generated cDNAs were converted to an Illumina Sequencing compatible RNA library using a Nextera XT DNA Library Preparation Kit (Illumina). Library concentration and quality were assessed using the Qubit 1X dsDNA HS Assay kit (Thermo Fisher Scientific) and the 4200 TapeStation with High Sensitivity D5000 ScreenTape Assay (Agilent Technologies). RNA libraries underwent partial lane sequencing on an Illumina Novaseq X platform (Illumina), with a sequence depth of 6 G and a read length of 2 x 150 bp. Sequencing was performed by Novogene AIT Genomics.

Raw sequencing read quality was assessed using FastQC (v0.12.1). Adapter removal and quality trimming were performed with fastp (v0.23.4) using --trim_poly_g, --trim_poly_x, -q 20, and -l 20. Trimmed reads were aligned to the human reference genome (hg38) using HISAT2 (v2.2.1). Gene-level read counts were quantified using Subread/featureCounts (v2.0.6) with the Ensembl GRCh38 release 114 annotation to generate a gene-by-sample count matrix. Ensembl gene IDs were converted to gene symbols using biomaRt (v2.58.0), and gene symbols were appended to the count matrix. TPM values were subsequently calculated from the raw counts.

For PCA, genes with low expression (total counts ≤ 10) were removed, and a variance stabilizing transformation (VST) was applied using DESeq2. PCA was performed on the transformed data using the prcomp function in R, and the first two principal components were visualized using ggplot2.

Differential expression analysis was performed using the voom–limma pipeline implemented in the limma and edgeR packages. Briefly, count data were normalized, transformed using voom, and fitted to a linear model. Correlation between replicate samples was accounted for using duplicateCorrelation. Differentially expressed genes were identified using empirical Bayes moderation, and p-values were adjusted using the Benjamini–Hochberg method.

For GSEA, genes were ranked based on the moderated t-statistics from the differential expression analysis. Hallmark gene sets (H) were obtained from the MSigDB using the msigdbr package. Enrichment analysis was performed using the clusterProfiler package with the fgsea algorithm. Gene sets with adjusted p-values < 0.05 were considered significantly enriched.

To compare pathway enrichment profiles, gene set enrichment results obtained in this study were compared with publicly available datasets, including active DTCs reported by Aouad et al.^21^ and reawakened DTCs from GSE289726^31^. For each dataset, enrichment results were matched based on common Hallmark gene sets. In comparison with the Aouad et al. dataset^21^, normalized enrichment scores (NES) from this study were compared with reported T-statistics for pathway activation. In the comparison with GSE289726, NES values derived from reanalysis of the dataset were directly compared with those obtained in this study. Significantly enriched pathways were defined using an adjusted p-value threshold of 0.05. Overlap between enriched pathways was assessed by classifying pathways based on significance and directionality, and the statistical significance of overlap was evaluated using Fisher’s exact test. Scatter plots were generated to visualize the relationship between enrichment scores across datasets, with point size reflecting statistical significance.

All analyses were performed in R (version 4.4.0).

### GSEA (desktop version)

GSEA was performed using GSEA v4.4.0 software (Broad Institute, http://www.broadinstitute.org/gsea/msigdb/index.jsp). For the desktop-based analysis, the number of permutations was set to 1000. Low-expressed genes, rRNAs, and mtRNAs were excluded from the input gene lists prior to analysis.

### RNAscope

RNAscope in situ hybridization (ISH) of FFPE samples was performed using the RNAscope 2.5 HD Reagent Kit-Brown (DAB-HRP) from Advanced Cell Diagnostics (ACD). Sample pretreatment and RNAscope were performed according to the manufacturer’s instructions (ACD). RNAscope probes, Hs-IFITM3-O1-C1 (Cat. No.1062531-C1) were purchased from ACD. Microscopic images were obtained using a NanoZoomer S210 digital slide scanner (Hamamatsu Photonics K.K.)

## Data availability

The RNA-seq datasets have been deposited in the Gene Expression Omnibus (RRID: SCR_005012) with the accession number GSEXXXXXX.

## Acknowledgements

We thank H. Kamioka for secretarial support. This work was supported by a JSPS Grant-in-Aid for Challenging Research (Pioneering) under Grant Number JP21K18259 (T.S.), a Takeda Science Foundation Research (T.S.), Project Mirai Cancer Research Grants (T.S.), the G-7 Scholarship Foundation (T.S.), Research Grant of the Princess Takamatsu Cancer Research Fund (T.S.), The Naito Foundation (T.S.), and Multilayered Stress Diseases (T.O.), and Medical Research Center Initiative for High Depth Omics (T.O.).

## Author information Contributions

K.H. and T.S. contributed equally to this work. T.S. conceived the study, K.H., T.S., Y.F., Y.M and H.Y. performed experiments and analyzed data. K.H., T.S. and T.O. wrote the manuscript. K.H. and T.S. performed bioinformatic analyses. N.I. provided experimental advice and discussion. H.H. provided cancer samples, advice and discussion. T.O. supervised the overall project.

## Ethics declarations Competing interest

The authors declare no competing interests.

