## Supplementary Figures 1-11, Supplementary Tables 1-2 for "Modeling the MRD state reveals the insomnia of chemotherapy-tolerant persister clones"

**Supplementary Fig 1. Hata et al.**

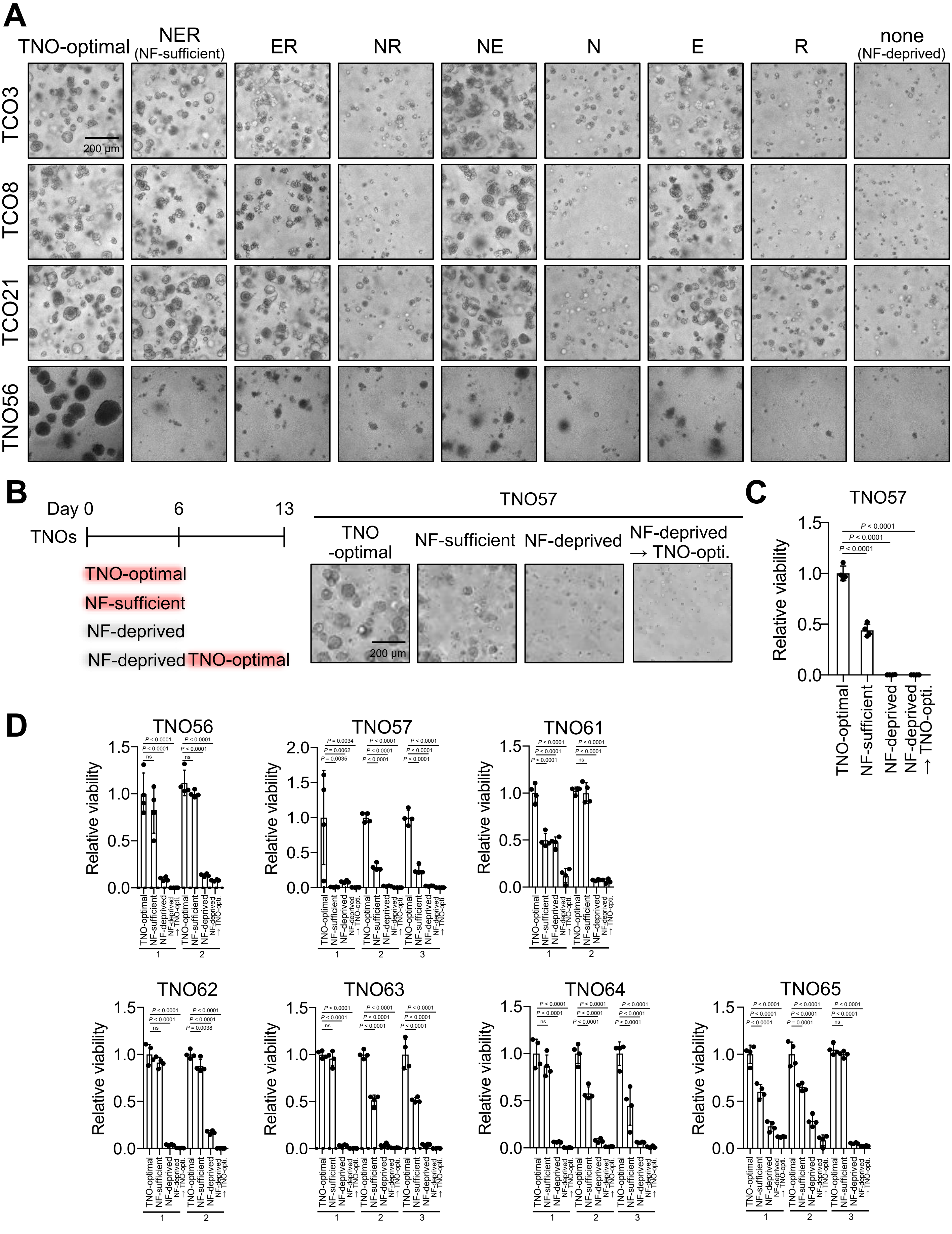

**Supplementary Fig 1 (Continued). Hata et al.**

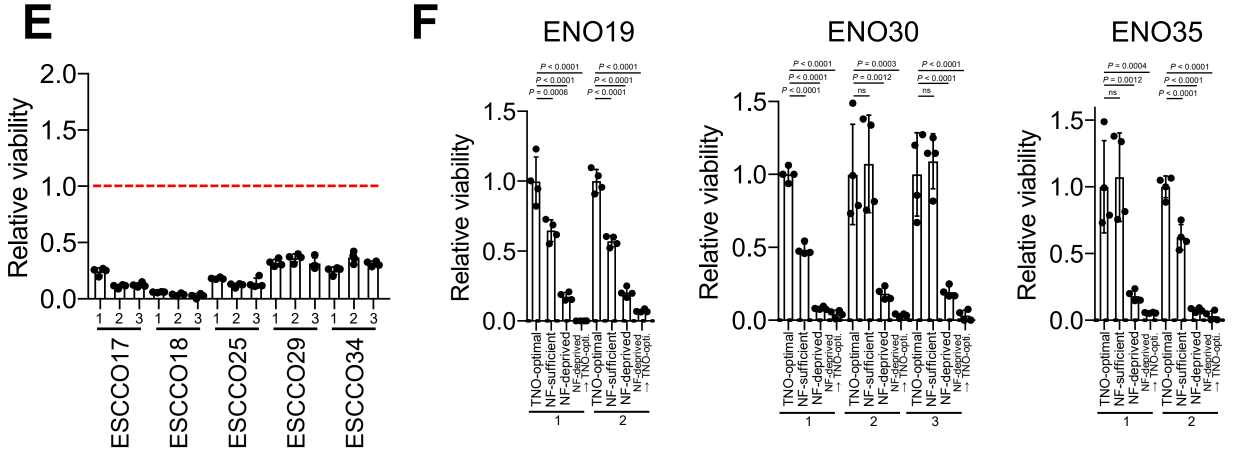


**Supplementary Figure 1. (related to Figure 2) Assessment of NF dependency in the organoid library**

**A,** Representative bright-field images of organoids cultured under different combinations of NFs at day 6. N: Noggin, E: EGF, R: R-spondin 1. Scale bar, 200 μm. **B, C,** Schedule for assessing NF dependency and the transition from NF-deprived to TNO-optimal, with representative bright-field images at day 6 (TNO-optimal, NF-sufficient, and NF-deprived) and day 13 (after transition from NF-deprived to TNO-optimal conditions) (**B**). Relative viability was calculated as the ratio of each value to that of TNO-optimal cultures (**C**). Statistical significance was determined by one-way ANOVA followed by Dunnett's multiple comparisons test. Scale bar, 200 μm. **D,** NF dependency and cell viability following the transition from NF-deprived to TNO-optimal conditions in additional TNOs. Statistical significance was determined by one-way ANOVA followed by Dunnett's multiple comparisons test. **E,** NF dependency in ESCCOs. Relative viability was calculated as the ratio of NF-deprived to NF-sufficient cultures. Data are presented as means ± SD from four independent experiments. ESCCO: esophageal squamous cell carcinoma organoid. **F,** NF dependency and cell viability following the transition from NF-deprived to TNO-optimal in ENOs. Statistical significance was determined by one-way ANOVA followed by Dunnett's multiple comparisons test. ENO: esophageal normal epithelial organoid.

### Supplementary Fig 2. Hata et al.

**A**

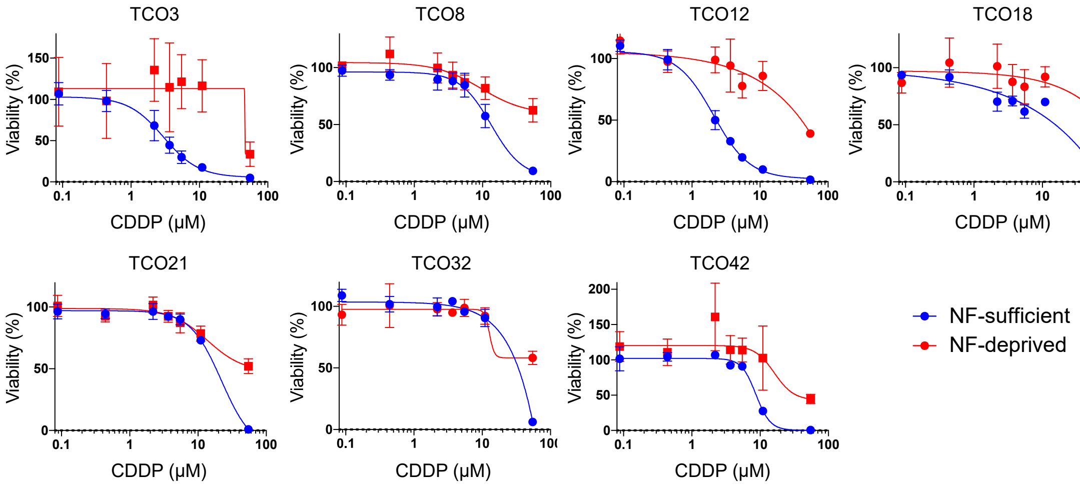


| IC50 (μM) | TCO3 | TCO8 | TCO12 | TCO18 | TCO21 | TCO32 | TCO42 |
|---|---|---|---|---|---|---|---|
| NF-sufficient | 2.999 | 13.43 | 2.077 | 12.80 | 22.06 | 32.12 | 8.679 |
| NF-deprived | 46.28 | >55 | >55 | >55 | >55 | >55 | >55 |

**B**

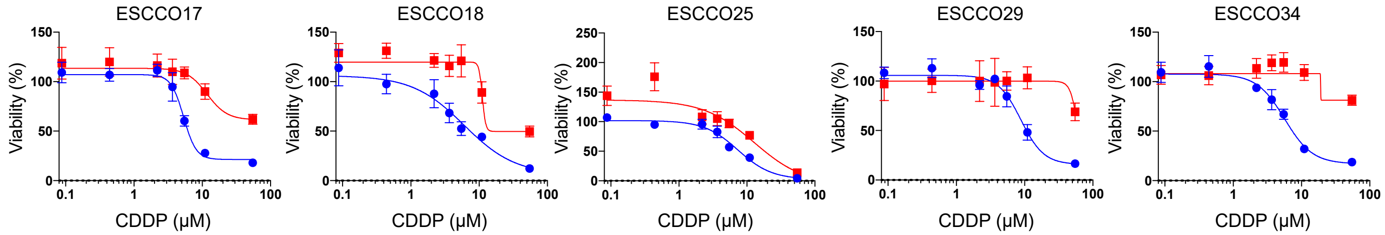


| IC50 (μM) | ESCCO17 | ESCCO18 | ESCCO25 | ESCCO29 | ESCCO34 |
|---|---|---|---|---|---|
| NF-sufficient | 5.338 | 6.011 | 2.448 | 8.844 | 5.673 |
| NF-deprived | >55 | >55 | 5.398 | >55 | >55 |

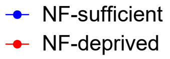


**Supplementary Figure 2. (related to Figure 2) Effects of NF deprivation on chemotherapy responses in tumor cells**

**A,** Dose-response curves for TCOs treated with CDDP under NF-sufficient and NF-deprived conditions. Cells were seeded on day 0, treated with CDDP from day 4 to day 7, and viability was measured on day 10 using CellTiter-Glo. Data are presented as means ± SD from four technical replicates. The IC50 values are also shown. Similar results were obtained in two independent experiments. **B,** Dose-response curves for ESCCOs treated with CDDP under NF-sufficient or NF-deprived conditions. Data are presented as means ± SD from four technical replicates. The IC50 values are also shown.

**Supplementary Fig 3. Hata et al.**

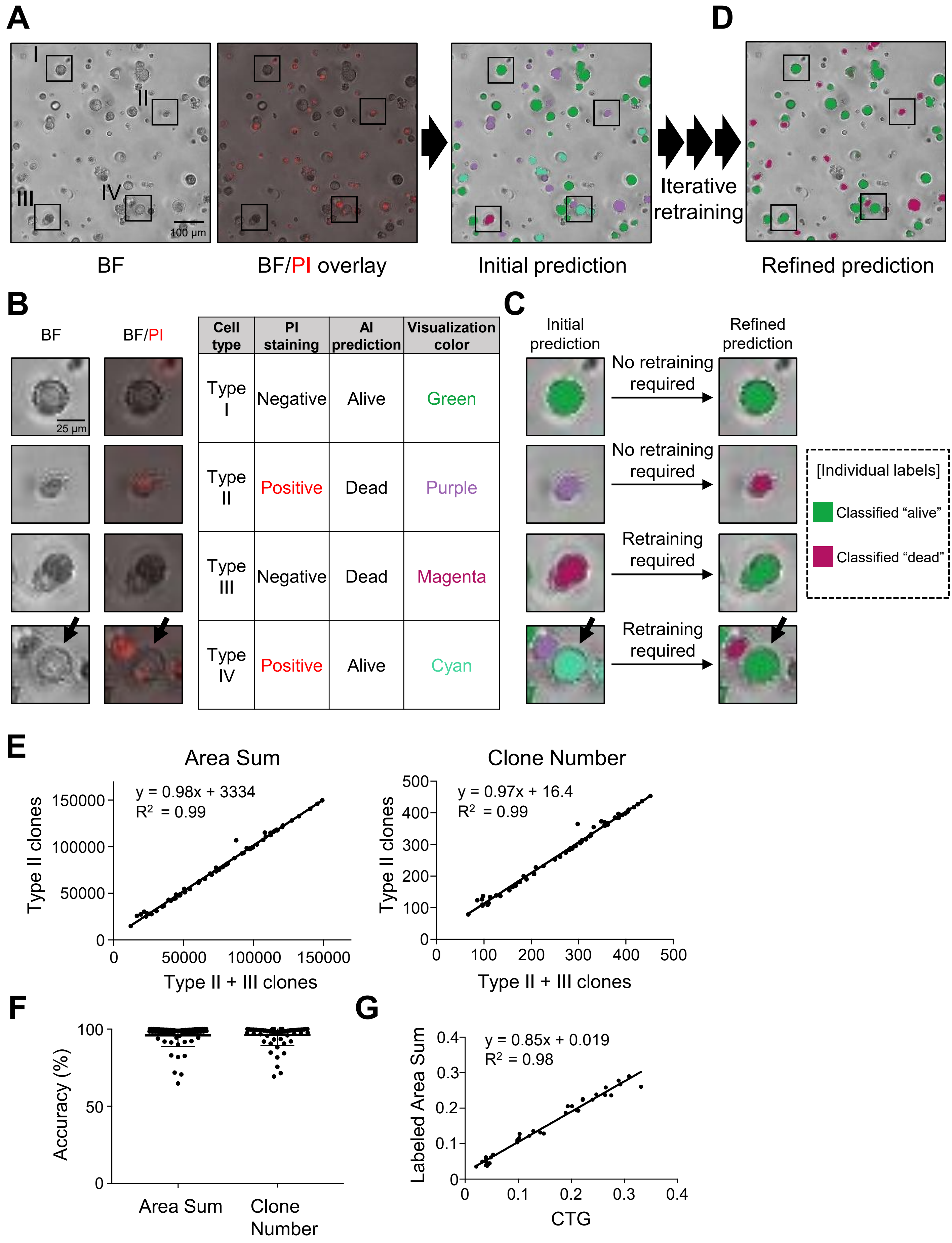


| Cell type | PI staining | AI prediction | Visualization color |
|---|---|---|---|
| Type I | Negative | Alive | Green |
| Type II | Positive | Dead | Purple |
| Type III | Negative | Dead | Magenta |
| Type IV | Positive | Alive | Cyan |

**Supplementary Figure 3. (related to Figure 2) Deep learning–based quantification of viable organoid clones from bright-field images**

**A-D,** Workflow of deep learning using paired bright-field and corresponding propidium iodide (PI)-stained images to infer live and dead clones from bright-field images alone. A deep learning model was trained to classify organoids as viable or dead based on bright-field images, using PI staining as ground truth (**A**). The trained model assigns pseudo-colors to each organoid as an initial prediction, and classifies clones into four types (Types I-IV) based on their positive/negative status for PI staining and the results of AI initial predictions. Type I: clones labeled as "alive" with negative PI staining, Type II: clones labeled as "dead" with positive PI staining, Type III: clones labeled as "dead" despite being PI-negative, Type IV: clones labeled as "alive" despite being PI-positive (**B**). Types I and II were excluded from additional training. In contrast, Type III clones were retrained as "alive" for additional training. Type IV clones (indicated by arrows) were basically reintroduced into the training dataset with confirmed annotations and were used for subsequent training, while morphologically obvious dead cells and debris were manually re-annotated (**C**). As a result of retraining, images with refined predictions were obtained (**D**). BF: Bright-field image. **E,** Correlation between PI-positive clones predicted as "Dead" (Type II) and all clones predicted as "Dead" (Type II and III clones) for total area of AI-annotated dead regions (Area Sum, left) and clone number (right) by linear regression analysis ($n = 66$). **F,** Accuracy of clonal labeling after deep learning. Accuracy was defined as the percentage of PI-positive clones predicted as "Dead" (Type II) among all clones predicted as "Dead" (Type II and III clones). Accuracy was 96.0 ± 7.2% for area sum, and 96.2 ± 6.6% for clone number. **G,** Correlation between cell viability measured by CellTiter-Glo (CTG) and total clonal area labeled by AI after deep learning.

**Supplementary Fig 4. Hata et al.**

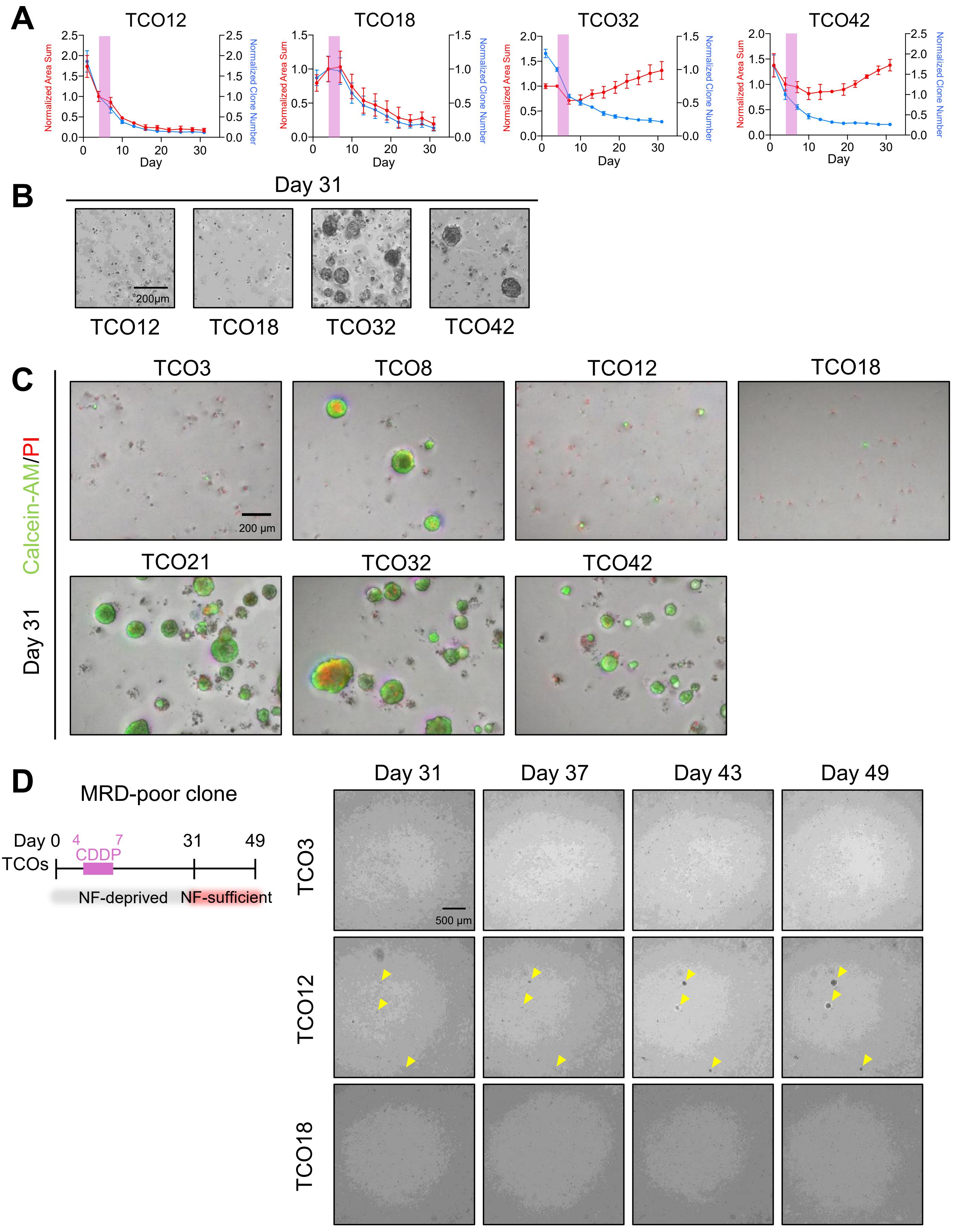

Supplementary Fig 4 (Continued). Hata et al.

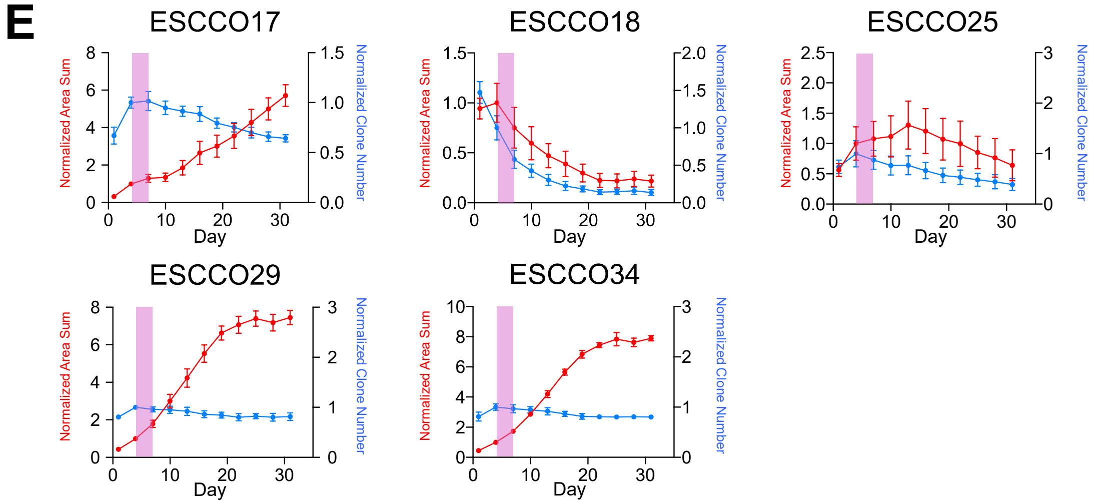


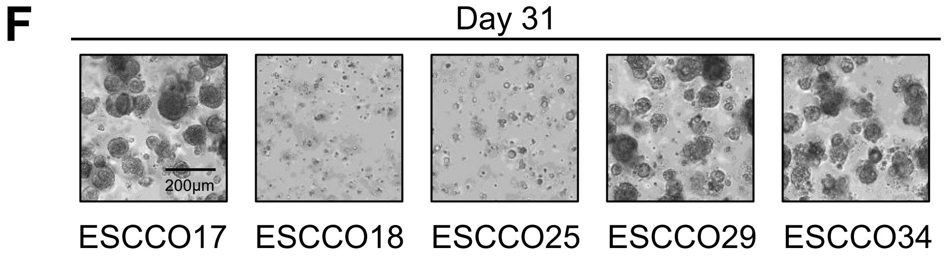


**Supplementary Figure 4. (related to Figure 2) Time-course analysis of tumor cells under NF-deprived conditions with chemotherapy treatment across TCO and ESCCO lines**

**A,** Time-course analysis of total clonal areas and clone numbers in cultures under NF-deprived conditions with concurrent chemotherapy exposure. Values were quantified from bright-field images and were normalized to day 4. The shaded regions indicate the period of CDDP treatment. **B,** Representative bright-field images from (**A**) on day 31. Scale bar, 200 µm. **C,** Each TCO line was cultured under NF-deprived conditions with concurrent chemotherapy exposure, as in **Fig. 2I**, and subsequently stained with Calcein-AM (green, live cells) and PI (red, dead cells). Representative images at day 31 are shown. Scale bar, 200 µm. **D,** Schedule of the transition from NF-deprived conditions with concurrent chemotherapy exposure to NF-sufficient conditions, with representative bright-field images from day 31 to day 49 (18 days after replacement with NF-sufficient medium). In representative images of TCO12, yellow arrowheads indicate expanding clones, which are extremely rare. Scale bar, 500 µm. **E,** Time-course analysis of total clonal areas and clone numbers in ESCCO cultures under NF-deprived conditions with concurrent chemotherapy exposure. Values were quantified from bright-field images and normalized to day 4. The shaded regions indicate the period of CDDP treatment. **F,** Representative bright-field images from (**E**) on day 31. Scale bar, 200 µm.

### Supplementary Fig 5. Hata et al.

A

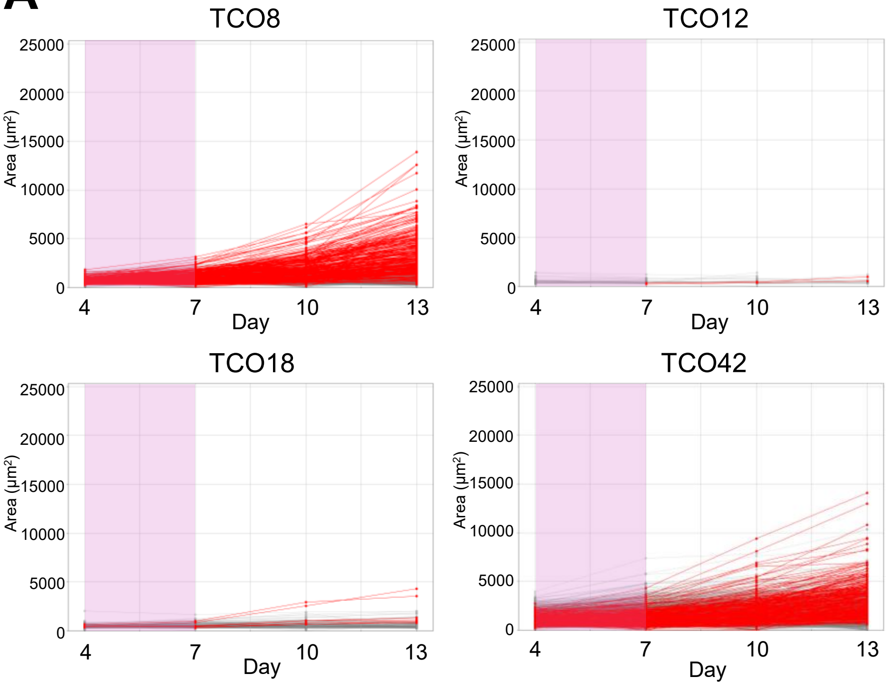


B

| | Total track number (clone) | Percentage of fusion (%) | Percentage of separation (%) |
|---|---|---|---|
| TCO3 | 843 | 0.5 (4/843) | 0.4 (3/843) |
| TCO8 | 1877 | 0.9 (16/1877) | 0.6 (11/1877) |
| TCO12 | 190 | 0.0 (0/190) | 0.0 (0/190) |
| TCO18 | 774 | 0.3 (2/774) | 0.3 (2/774) |
| TCO21 | 2455 | 1.5 (36/2455) | 1.4 (34/2455) |
| TCO32 | 1667 | 1.3 (22/1667) | 1.4 (23/1667) |
| TCO42 | 2443 | 1.6 (39/2443) | 2.0 (48/2443) |

**Supplementary Figure 5. (related to Figure 3) Longitudinal clone tracking under NF-deprived conditions with CDDP treatment**

**A,** Changes in clonal areas of individual clones from day 4 to day 13, quantified from bright-field images. Red lines indicate CP clones (≥ 2-fold expansion in area between day 7 and day 13), whereas gray colored lines indicate non-CP or dead clones (< 2-fold change). The shaded regions indicate the period of CDDP treatment. **B,** Table summarizing the total number of tracked clones, along with the percentage and number of clones undergoing fusion or separation.

**Supplementary Fig 6. Hata et al.**

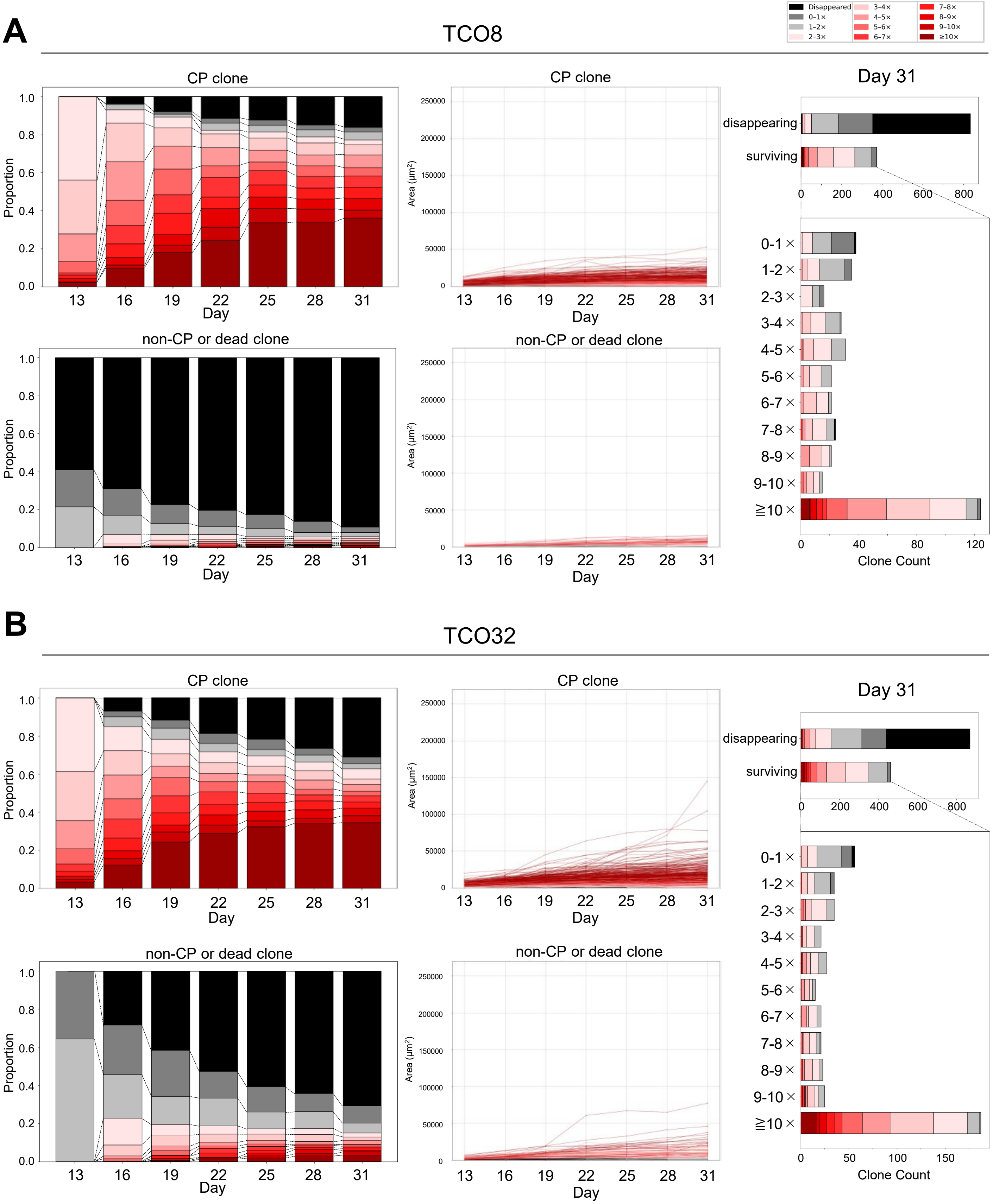

Supplementary Fig 6(Continued). Hata et al.

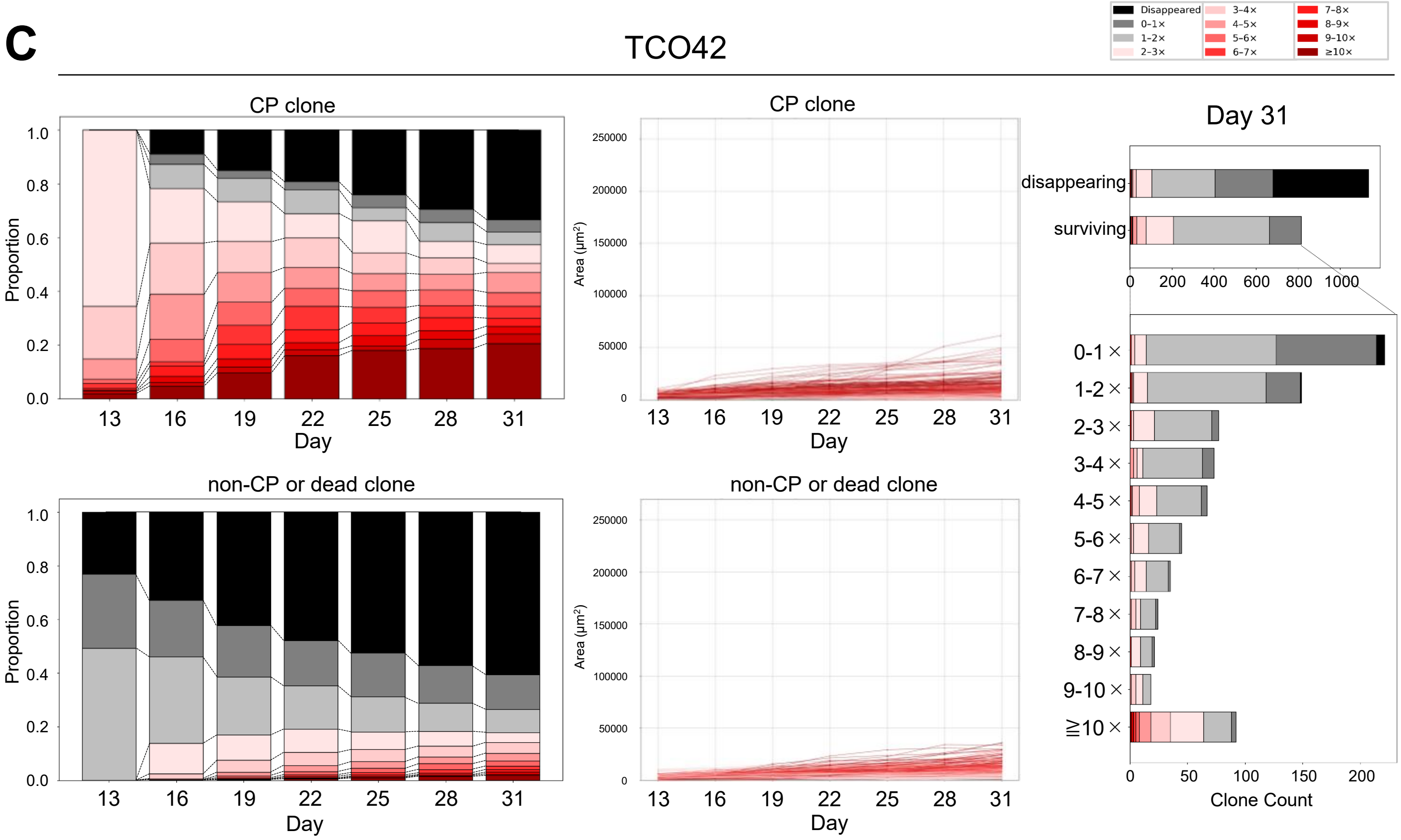


**Supplementary Figure 6. (related to Figure 3) Clone tracking in CP-rich TCO lines reveals clonal dynamics under chemotherapy treatment**

**A,** (left upper) 100% stacked bar graph showing the distribution of fold changes in clonal areas of CP clones of TCO8 at each time point (days 13-31). Fold changes were calculated for each clone at each time point relative to day 7 and were categorized into discrete ranges, with increasing values shown as progressively darker shades of red (≥10-fold as the darkest), 1-2-fold in light gray, and 0-1-fold in dark gray, and disappeared clones in black. Notably, only clones exhibiting ≥ 10-fold expansion relative to day 7 progressively become dominant over time, suggesting that a limited subset of clones possesses sustained proliferative capacity and may ultimately contribute to tumor outgrowth. (middle upper) Changes in the clonal areas of individual CP clones from day 13 to day 31. The color of each line corresponds to the categories shown in **Fig. 3D**. (left lower) 100% stacked bar graph showing the distribution of fold changes in clonal areas for non-CP or dead clones of TCO8 at each time point (days 13-31). Fold changes were calculated for each clone at each time point relative to day 7 and were categorized into discrete ranges, with increasing values shown as progressively darker shades of red (≥ 10-fold as the darkest), 1-2-fold in light gray, and 0-1-fold in dark gray, and disappeared clones in black. Notably, few non-CP clones at day 13 become expanding clones over time. (middle lower) Changes in the clonal areas of individual non-CP or dead clones from day 13 to day 31 (left). The color of each line corresponds to the categories shown in **Fig. 3F**. (right) Tracked clones were first categorized based on whether they survived or disappeared by day 31 (upper). From another perspective, surviving clones were further stratified based on changes in clonal area at day 31 relative to day 7 (lower). Stacked bar graphs show changes in the clonal areas of individual clones at day 13 relative to day 7. The color of each segment corresponds to the categories shown in **Fig. 3D, F**. **B,** The same clone tracking analysis described in **Supplementary Fig. 6A**, was applied to TCO32. **C,** The same clone tracking analysis described in **Supplementary Fig. 6A**, applied to TCO42.

#### Supplementary Fig 7. Hata et al.

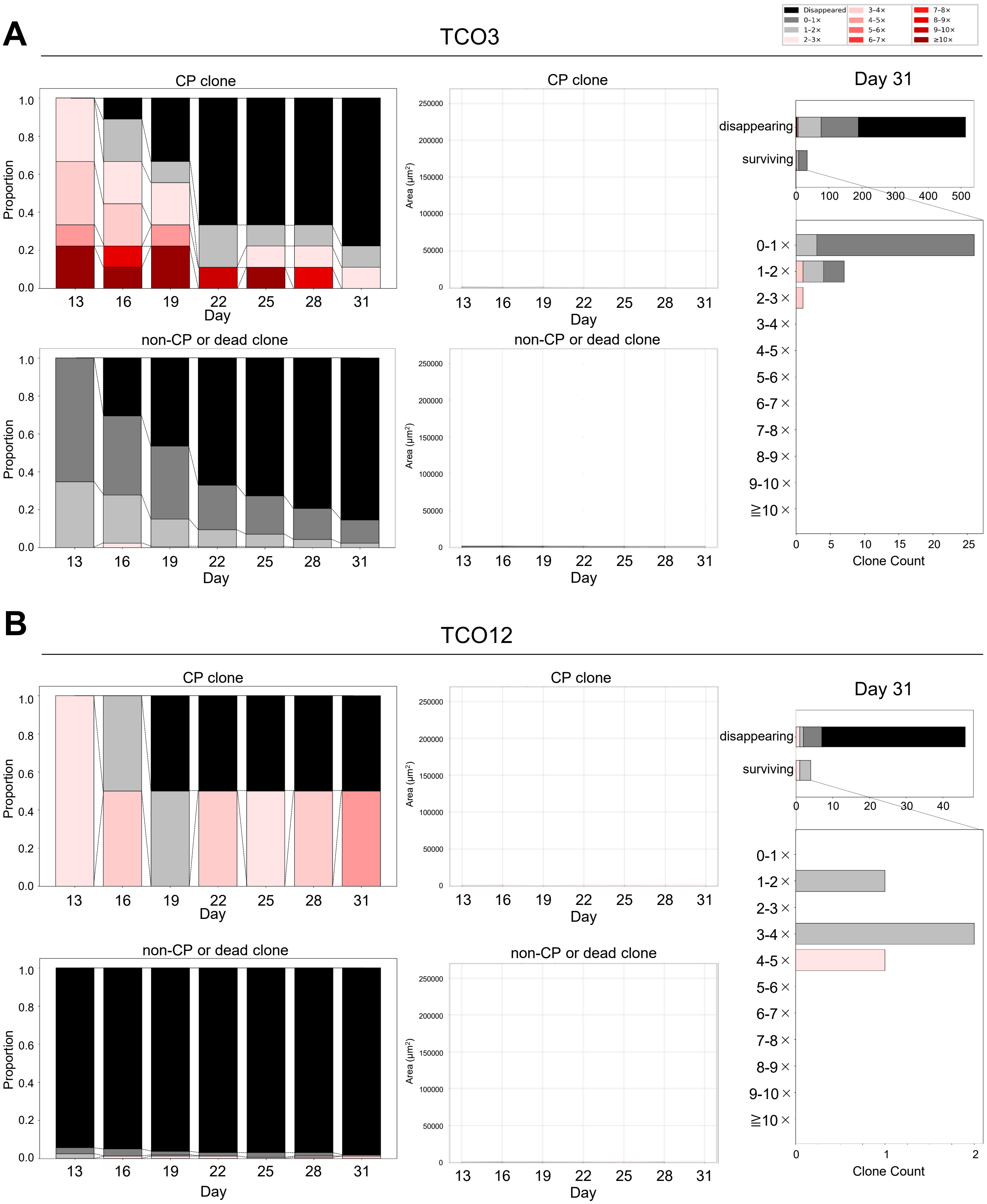

**Supplementary Fig 7(Continued). Hata et al.**

**C**

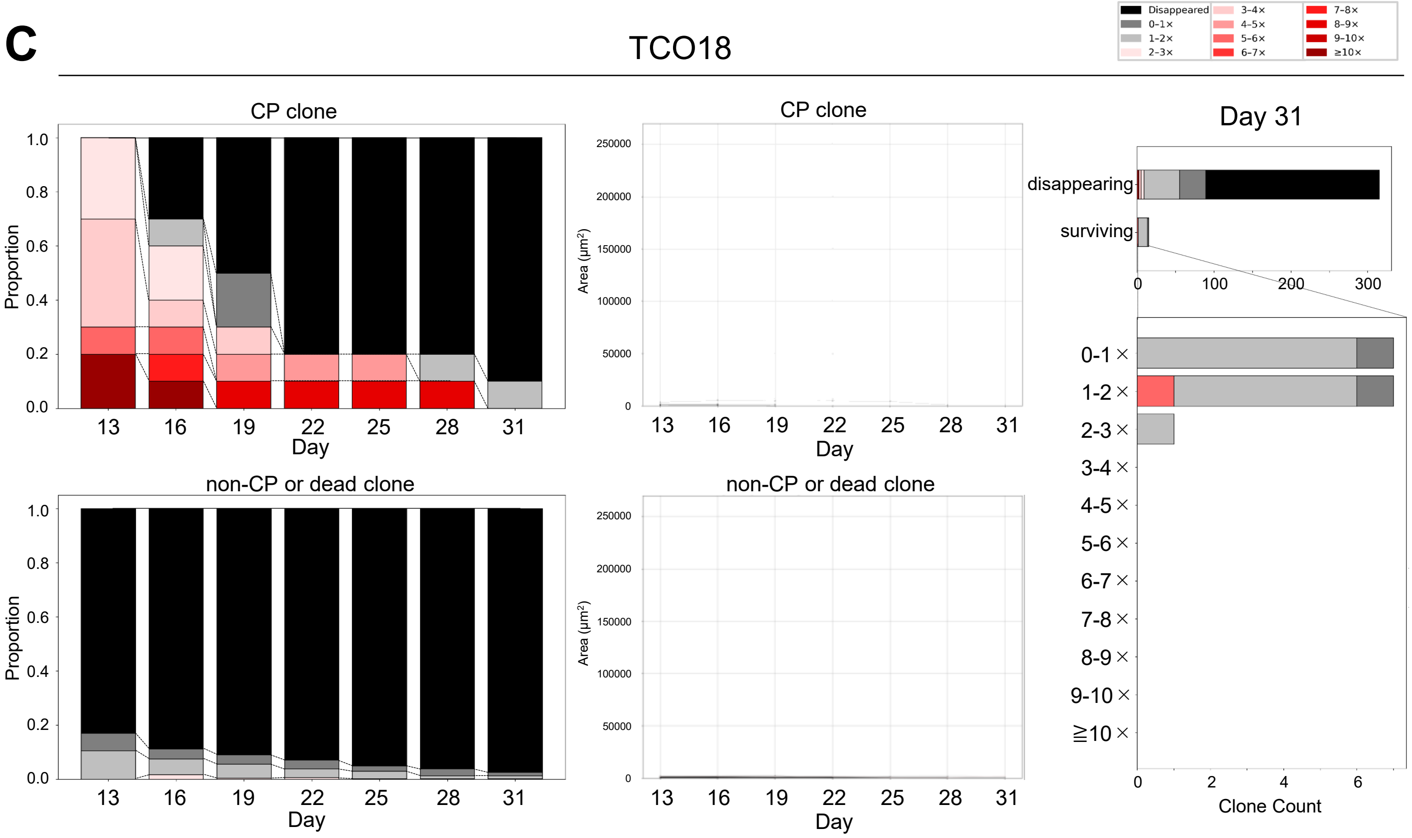


**Supplementary Figure 7. (related to Figure 3) Clone tracking in CP-poor TCO lines reveals limited clonal persistence and re-expansion under chemotherapy**

**A,** (left upper) 100% stacked bar graph showing the distribution of fold changes in clonal areas for CP clones of TCO3 at each time point (days 13-31). Fold changes were calculated for each clone at each time point relative to day 7 and were categorized into discrete ranges, with increasing values shown as progressively darker shades of red (≥ 10-fold as the darkest), 1-2-fold in light gray, and 0-1-fold in dark gray, and disappeared clones in black. (middle upper) Changes in clonal areas of individual CP clones from day 13 to day 31. The color of each line corresponds to the categories shown in **Fig. 3D**. (left lower) 100% stacked bar graph showing the distribution of fold changes in clonal area for non-CP or dead clones of TCO3 at each time point (days 13-31). Fold changes were calculated for each clone at each time point relative to day 7 and were categorized into discrete ranges, with increasing values shown as progressively darker shades of red (≥ 10-fold as the darkest), 1-2-fold in light gray, and 0-1-fold in dark gray, and disappeared clones in black. Notably, few non-CP clones at day 13 become expanding clones over time. (middle lower) Changes in the clonal areas of individual non-CP or dead clones from day 13 to day 31 (left). The color of each line corresponds to the categories shown in **Fig. 3F**. (right) Tracked clones were first categorized based on whether they survived or disappeared by day 31 (upper). From another perspective, surviving clones were further stratified based on changes in clonal areas at day 31 relative to day 7 (lower). Stacked bar graphs show changes in the clonal areas of individual clones at day 13 relative to day 7. The color of each segment corresponds to the categories shown in **Fig. 3D, F**. **B,** The same clone tracking analysis described in **Supplementary Fig. 7A**, applied to TCO12. **C,** The same clone tracking analysis described in **Supplementary Fig. 7A**, was applied to TCO18.

**Supplementary Fig 8. Hata et al.**

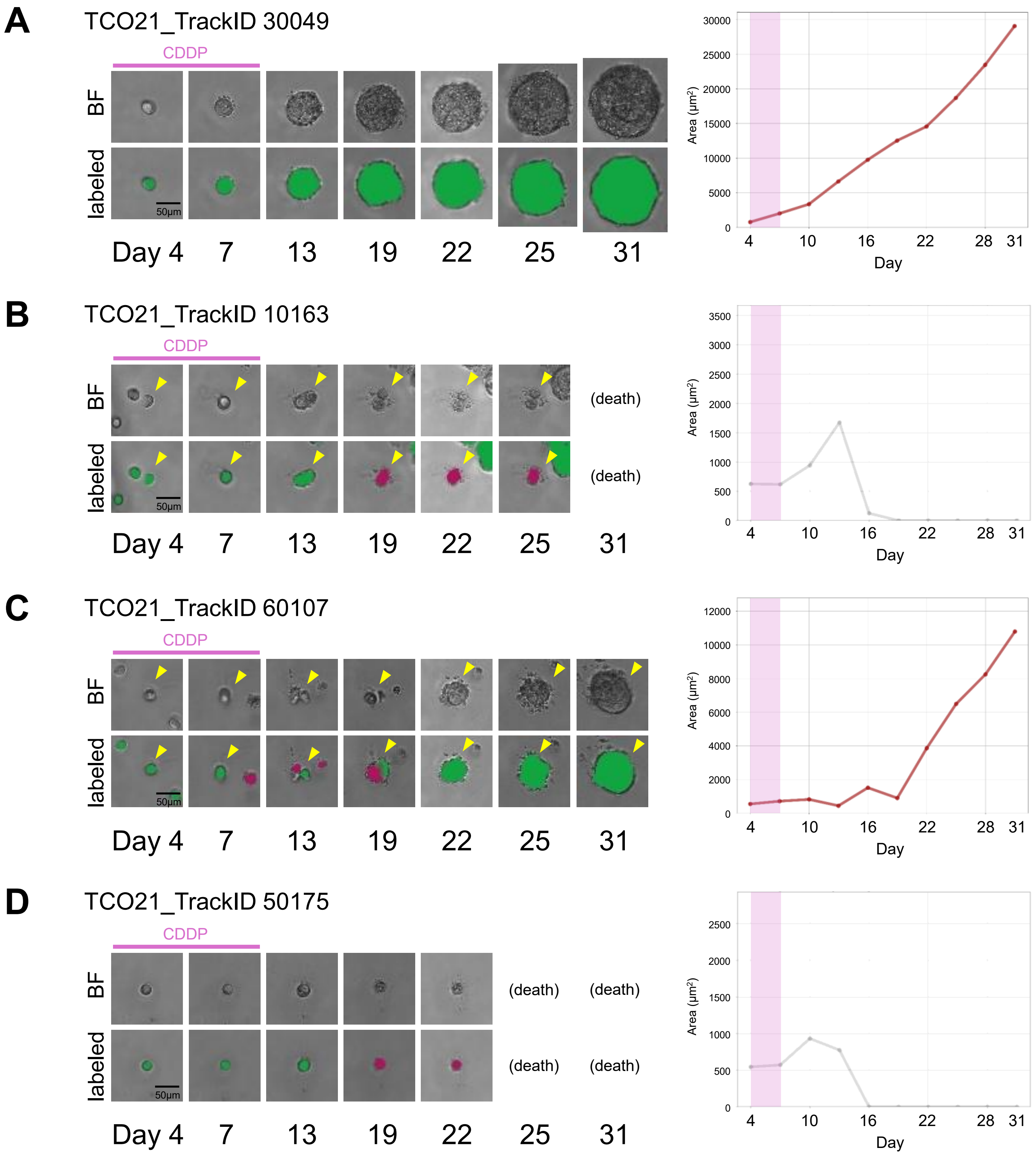

**Supplementary Figure 8. (related to Figure 3) Representative tumor clones exhibiting diverse longitudinal growth patterns**

**A,** Representative bright-field images and corresponding labeled images at days 4, 7, 13, 19, 22, 25, and 31 for a CP clone that ultimately expanded (left). Changes in the clonal area of the clone from day 4 to day 31 are shown (right). The shaded region indicates the period of CDDP treatment. **B,** Representative bright-field images and corresponding labeled images (left), along with a time-course plot of clonal area (right), for a CP clone that ultimately disappeared. The arrowheads indicate the tracked clone. **C,** Representative bright-field images and corresponding labeled images (left), along with a time-course plot of clonal area (right), for a non-CP clone that ultimately expanded. The arrowheads indicate the tracked clone. **D,** Representative bright-field images and corresponding labeled images (left), along with a time-course plot of clonal area (right), for a non-CP clone that ultimately disappeared. Scale bars, 50 μm.

### Supplementary Fig 9. Hata et al.

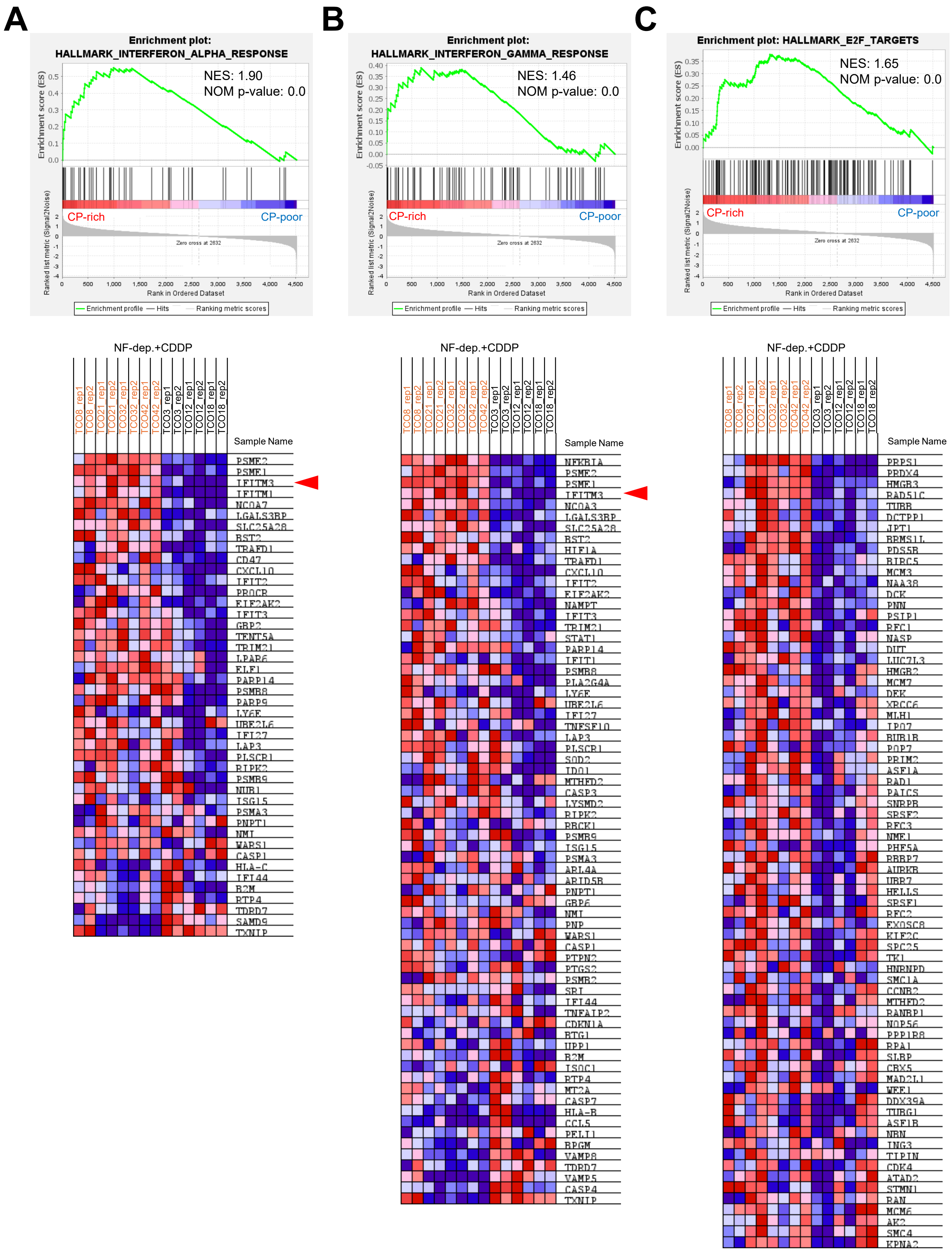

**Supplementary Figure 9. (related to Figure 4) Overrepresented pathways in CP-rich TCO lines identified by GSEA**

**A-C,** GSEA plots showing a comparison of transcriptional profiles between CP-rich and CP-poor TCO lines. Corresponding GSEA enrichment plots (upper) and heatmaps of the associated leading-edge genes (lower) for the top three significantly enriched pathways ranked by NES are shown, including HALLMARK_INTERFERON_ALPHA_RESPONSE (**A**), HALLMARK_INTERFERON_GAMMA_RESPONSE (**B**), and HALLMARK_E2F_TARGETS (**C**). Red arrowheads indicate *IFITM3*.

Supplementary Fig 10. Hata et al.

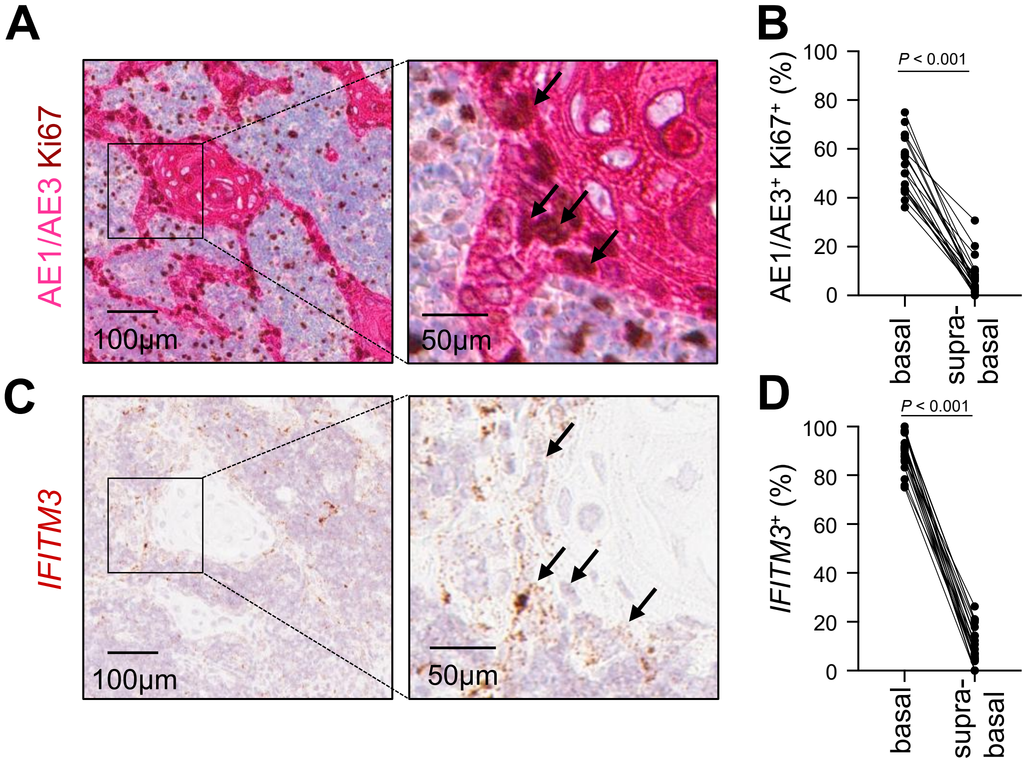


**Supplementary Figure 10. (related to Figure 4) High *IFITM3* expression in proliferating tumor cells in metastatic lymph nodes**

**A-C,** Immunostaining of Ki67 with AE1/AE3 and detection of *IFITM3* mRNA by RNAscope in another pathological specimen from Patient 5 in **Fig. 1C**. Ki67-positive with AE1/AE3-positive cells and *IFITM3*-positive cells were counted in two defined regions (basal cells and supra-basal cells). Representative double immunostaining for Ki67 and AE1/AE3 (**A**), and RNAscope-based detection of *IFITM3* mRNA (**C**) in serial pathological sections from Patient 5 shown in **Fig. 1C**. Corresponding cells positive for both markers in serial sections are indicated by arrows. The frequencies of Ki67- and AE1/AE3-double positive proliferating tumor cells (**B**), and *IFITM3*-expressing cells (**D**), were evaluated in the basal and supra-basal layers of the lymph node tumor nests. Results of immunostaining and RNAscope were reviewed and confirmed by a pathologist. Statistical significance was determined by a two-sided Wilcoxon test. Scale bars, 100 µm (left panels of **A** and **C**), 50 µm (right panels of **A** and **C**).

### Supplementary Fig 11. Hata et al.

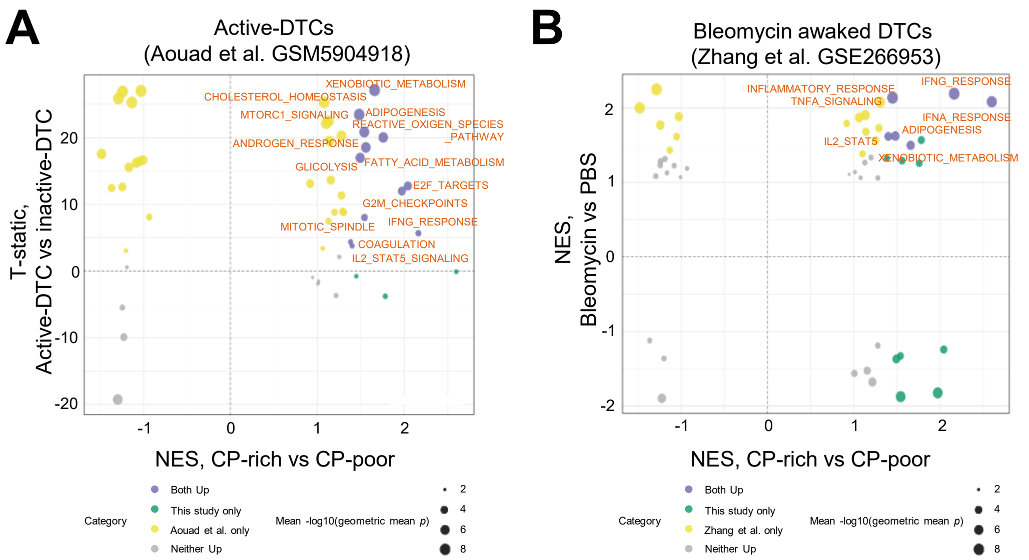


**Supplementary Figure 11. (related to Figure 4) The transcriptional similarity between our CP population and persister-like cells identified in previously reported models.**

**A**, Identification of commonly enriched pathways between TCO-derived CPs and active DTCs reported by Aouad et al.[21] in a mouse model of lung metastatic breast cancer. **B**, Identification of commonly enriched pathways between TCO-derived CPs and reawakened DTCs reported by Zhang et al.[31] in a bleomycin-induced lung inflammation model.

#### Supplementary Table 1. Hata et al.

| Tongue cancer Patient no. | Sex | Age | Cancer location | Type | pTNM | Stage | Organoid formation | | | Source |
|---|---|---|---|---|---|---|---|---|---|---|
| | | | | | | | Normal | Cancer | Lymph node | |
| 3 | F | 85 | tongue | SCC | T4aN0M0 | IVA | Yes | Yes | N/A | Sase et al., 2025 |
| 8 | F | 50 | tongue | SCC | T2N0M0 | II | Yes | Yes | N/A | Sase et al., 2025 |
| 12 | F | 77 | tongue | SCC | T2N0M0 | II | Yes | Yes | N/A | Sase et al., 2025 |
| 18 | M | 73 | tongue | SCC | T3N0M0 | III | Yes | Yes | N/A | Sase et al., 2025 |
| 21 | M | 85 | tongue | SCC | T2N3bM0 | IVB | Yes | Yes | N/A | Sase et al., 2025 |
| 32 | M | 91 | tongue | SCC | T2N0M0 | II | Yes | Yes | N/A | Sase et al., 2025 |
| 42 | F | 55 | tongue | SCC | T4aN0M0 | IVA | Yes | Yes | N/A | This study |
| 52 | M | 59 | tongue | SCC | T3N3bM0 | IVB | Yes | No | Yes | This study |
| 56 | F | 50 | tongue | SCC | T3N1M0 | III | Yes | No | N/A | This study |
| 57 | M | 63 | tongue | SCC | T4aN3bM0 | IVB | Yes | Yes | Yes | This study |
| 61 | M | 63 | tongue | SCC | T2N2bM0 | IVA | Yes | Yes | Yes | This study |
| 62 | M | 66 | tongue | SCC | T4aN1M0 | IVA | Yes | No | No | This study |
| 63 | M | 62 | tongue | SCC | T4aN0M0 | IVA | Yes | No | N/A | This study |
| 64 | M | 38 | tongue | SCC | T4aN3bM0 | IVB | Yes | Yes | No | This study |
| 65 | M | 38 | tongue | SCC | T3N2aM0 | IVA | Yes | Yes | N/A | This study |

N/A, not applicable.

#### Supplementary Table 2. Hata et al.

| Esophageal cancer Patient no. | Sex | Age | Cancer location | Type | pTNM | Stage | Organoid formation | | Source |
|---|---|---|---|---|---|---|---|---|---|
| | | | | | | | Normal | Cancer | |
| 17 | M | 81 | esophagus | SCC | T3N1M0 | IIIB | Yes | Yes | Nakagawa et al., 2025 |
| 18 | M | 65 | esophagus | SCC | T1bN0M0 | IB | Yes | Yes | Nakagawa et al., 2025 |
| 19 | M | 67 | esophagus | SCC | T3N1M0 | IIIB | Yes | Yes | Nakagawa et al., 2025 |
| 25 | M | 61 | esophagus | SCC | T2N2M0 | IIIB | Yes | Yes | Nakagawa et al., 2025 |
| 29 | M | 73 | esophagus | SCC | T3N1M0 | IIB | Yes | Yes | Nakagawa et al., 2025 |
| 30 | F | 80 | esophagus | SCC | T3N3M0 | IVA | No | Yes | Nakagawa et al., 2025 |
| 34 | M | 73 | esophagus | SCC | T3N1M0 | IIIB | Yes | Yes | Nakagawa et al., 2025 |
| 35 | M | 49 | esophagus | SCC | T3N2M0 | IIIB | Yes | Yes | Nakagawa et al., 2025 |

N/A, not applicable.
